# Neural Mass Model of Auditory stimulus responses in Marmoset Cortex

**DOI:** 10.1101/2025.07.27.665709

**Authors:** Bernard A Pailthorpe

## Abstract

A composite, heterogeneous neural mass model of the marmoset auditory and pre-frontal cortex (pFC) is used to model negativity mismatch experiments. This builds upon previous analysis of a compact pFC cluster. Simulated average deviant response delays were 132 ms for Auditory areas, and 147 ms for pFC areas, comparable to the 100-250 ms experimentally measured range. Details of responses for each anatomical area are presented. Area TPO is a key output node from the auditory cluster with strong links to pFC. Area AuA1 is activated by the deviant, high frequency, stimulus then turns off after the standard stimuli cease. A NMM response and delay is observed in the pFC sub cluster. Inactivation of AuRPB causes ambiguous spikes in AuA1 and reduced responses, consistent with experimental observations. The simulations facilitate analysis of the driving forces arriving at AuA1 and suggest that feedback from AuRPB to AuA1 is carried over multiple local network pathways. The simulations reproduce the experimentally observed delays and dissect the roles of participating areas and pathways.

## 1. Introduction

Negativity mismatch (NMM) experiments probe auditory responses and have been reported in a number of animal models. Extensive results are available for marmoset and are modelled herein. The simulations aim to provide a computational model of the NMM experiments (eg. Komatsu et. al. 2023; Obara et. al. 2023). The tonotopic map of marmoset auditory cortex responses (Tani et. al. 2018) identify the relative high and low frequency of each auditory area and inform the simulation protocol used herein to model the NMM experiments. A deviant, “high” frequency, stimulus pulse replaces a central pulse in a stream of 10 standard, “low” frequency, pulses – analogous to the oddball experimental paradigm (Obara et. al. 2023). Here a neural mass model and simulations of these experiments is presented that reproduces the observed delays in response, and highlights the role of individual areas.

## 2. Methods

Network methods and visualisation were used to identify a nine node cluster of Neural Masses (NM) comprising auditory areas, following my earlier analysis (Pailthorpe 2024a) of the marmoset connectivity data (Majka et. al. 2016, 2020; Rosa et. al. 2024). A 3D model of pre Frontal Cortex (pFC), comprising 6 NM was developed (Pailthorpe, 2024a), along with analysis of its dynamics and transitions (Pailthorpe, 2024b). Available data on those anatomical areas in pFC facilitated assignment of frequency bands and parameters to each NM, as listed in Table 1. That analysis is here extended to auditory anatomical areas. The local sub clusters studied are: the 9 auditory areas are: AuCPB, AuA1, AuRT, AuAL, TPO, AuML, AuCL, AuCM and AuRPB. Note that TPO, in the temporal lobes, is included since network analysis places it in the auditory module (Pailthorpe, 2024a) and it is closely linked to the other areas listed. For pFC the 8 areas are: A10, A32V, A9, A32, A46D, A11, A8aD and A8b. The strongest links present are within each sub cluster. The strongest link from auditory areas forward to pFC is from TPO (a temporal area) to A32V, and the strongest feedback is again between those two nodes. Other significant direct links forward are: AuCPB – A32V, and TPO – A11, -A46D, -A9 and -A8b. Areas A9 and A32 received only very weak direct links from any of the 9 auditory areas, yet are locally enmeshed with the other pFC areas so need to be included. Areas A8aD and A8b appear to be key output nodes of pFC, with medium strength (weights 228 and 340, respectively) on the Area 6DR, a premotor area, justifying their inclusion.

**Table 1.** Nodes of the sub-clusters in the model. Left: 9 auditory nodes. Right. 8 pFC nodes. Frequency bands found for the eight nodes in the pFC cluster. Neural mass parameters were tuned to the listed frequency band. The resultant dominant frequency f was found from peaks in the power spectral density of dy(t) for each node under constant stimulus (1 G noise + 100 Hz pulse rate to all nodes); secondary peaks are listed when significant.

| Node # | Area | Frequency band | f (Hz) |
| --- | --- | --- | --- |
| 1 | AuCPB | alpha/gamma | 11.5, 17.3 |
| 2 | AuA1 | theta-gamma | 17.5, 11.7, 5.8, 40.9 |
| 3 | AuRT | beta -gamma | 6.3, 12.6, 18.9 .. 44.3 |
| 4 | AuAL | alpha | 10.0 |
| 5 | TPO | gamma | 17.5, 35 |
| 6 | AuML | alpha | 16.1 |
| 7 | AuCL | theta | 5.44 |
| 8 | AuCM | beta | 9.94, 17.5 |
| 9 | AuRPB | alpha | 10.0 |
| 10 | A10 | gamma | 32.44 |
| 11 | A32V | theta | 5.06, 10.7 |
| 12 | A32 | beta | 16.0 |
| 13 | A9 | alpha | 9.7 |
| 14 | A46D | theta | 5.5 |
| 15 | A11 | beta | 15.2, 32.44 |
| 16 | A8aD | alpha | 9.9 |
| 17 | A8b | beta | 18.3 |

The simulated model uses 17 neural masses, 9 for the auditory cortex and 8 for pFC, interacting via the experimental link weights (Majka et. al. 2016; Rosa et. al. 2024, Pailthorpe 2024a). The model builds on the classical Wilson-Cowan (WC) and Jansen-Ritt (JR) methods, as extended in (Pailthorpe 2024b) for Marmoset pFC. That work incorporated local link weights into a. new form of the voltage – rate transformation function. The classic 3-population NM model did not produce the expected responses to stimuli of the auditory areas. It appears that a NM model capable of gamma bursting is needed – here I adopt the Wendling model comprising a mixed 5- and 3-population NMs, and including a fast inhibitory neural sub-population (Wendling et. al. 2002]) that was later re-parameterised [Ersoz et. al. 2020].

This successfully describes EEG patterns and transitions. Data for 3 anatomical areas – AuCPB, AuA1 and AuRT – were ambiguous in assigning characteristic frequencies for the NM’s, so those were modelled by 5 sub-populations and, hence, capable of gamma bursting. Clusters of 6, 8 or 9 auditory areas were explored. Ultimately 9 areas were used, providing coverage of core, belt and parabelt regions and including RPB (identified as a critical link in NMM response (Obara et. al. 2023). NMs 4, AuAl 6, AuML and 9, AuRPB correspond to areas that were not tracer injection sites, so there is no in-link data for those areas. Those weights were estimated from observed statistical trends in the available data (to be published). Simulations were run for 20 s, with stimuli applied at 4 s, well past transient dynamics.

Direct Auditory - pFC links are included naturally via the available marmoset connectivity matrix (Rosa et. al. 2024). It is likely that multiple neural pathways connect distant areas, with some 2-step links via intermediate waypoints providing longer travel distances and signal delays. In all 443 pathways, via up to 14 waypoints (located anterior to bregma, y > 0 mm, A-P axis) were used with composite pathway lengths up to 30.7 mm. Composite weights were calculated as the geometric mean of contributing weights. The intermediate areas include the temporal, cingulate and parietal, but not visual, areas. The majority of those links were of very small weight so contributed little stimulus. Important waypoints, with strong links, were TE3 (Temporal areas), A23b (Cingulate) and PGa-IPa in Posterior Parietal Cortex (PPC). Observations of a positive deviance response in parietal areas of marmoset (Komatsu et. al. 2023) lend credence to the inclusions of these waypoints. The connectivity data lists numerous, weaker 2-step links from auditory areas to pFC via Opt, PF and PG in PPC, with the strongest originating in TPO. Additionally brain-spanning neural fields (Robinson et. al. 2016) can provide a communication pathway been areas – that is not included here, since it would require more sophisticated simulations. Generally simulations using only direct links elicited weaker responses in pFC, so the 2-step pathways were included in the simulations reported below, following our earlier study of visual stimuli (Pailthorpe 2023). That may result in an overestimated response, since not all waypoints might participate. Some comparisons to the direct links only are provided in the Supplementary Information. Direct feedback links from pFC to auditory areas were generally weak, so no such 2-step links were explored.

The simulations aim to provide a computational model of the NMM experiments (Komatsu et. al. 2023; Obara et. al. 2023). Thus the difference between responses to two sets of stimuli were calculated. In a simple case, 10 smooth pulse functions (cf. Figure 1) were applied to the low frequency sensitive areas, being those identified by experiment (Tani et. al., 2018). Fractional weights of stimulus to each area was proportional to the measured (Ca) voltage response to low or high frequency tones, rescaled to a fraction: 0-1. That was compared to a deviant stimulus in which the middle (#5) pulse was replaced by a smooth pulse function directed to the high frequency sensitive areas (representing the experimental high frequency tone), as also identified (Tani et. al. 2018). Following Figs. 4, 5 and Table 1 of (Tani et. al. 2018) the low frequency responsive areas are: AuCPB NM#1 (fraction 0.6), AuA1 #2 (0.3), AuRT #3 AuRT (0.6), #4 AuAL (1.0), #6, AuML (0.4), AuRPB #9 (0.6), with weight fractions shown in brackets. The high frequency responsive areas are: AuCPB #1 (0.5), AuA1 #2 (0.5), AuML #6 (0.5), AuCL #7 (1) and AuRPB #9 (0.7). To be explicit: four areas AuCPB, AuA1, AuML and AuRT are common to the standard and deviant stimuli. In the standard stimulus (low f) areas AuRT and AuAL, are added; while for the deviant (high f) stimulus AuCL is included. The responses in those areas are likely to provide hints as to the origin and mechanisms of the deviant response. Signalling delays are incorporated as time delays in the model due to transmissions at axon velocity, here assumed to be 1 m/s (unmyelinated) locally within each cluster (≅ 2 - 5 mm links) and also for direct links between the two clusters, with lengths 10 – 16 mm. For the longer range 2-step links between the two clusters (lengths ≅ 12-30 mm) transmission velocity was taken to be 10 m/s.

**Figure 1.**
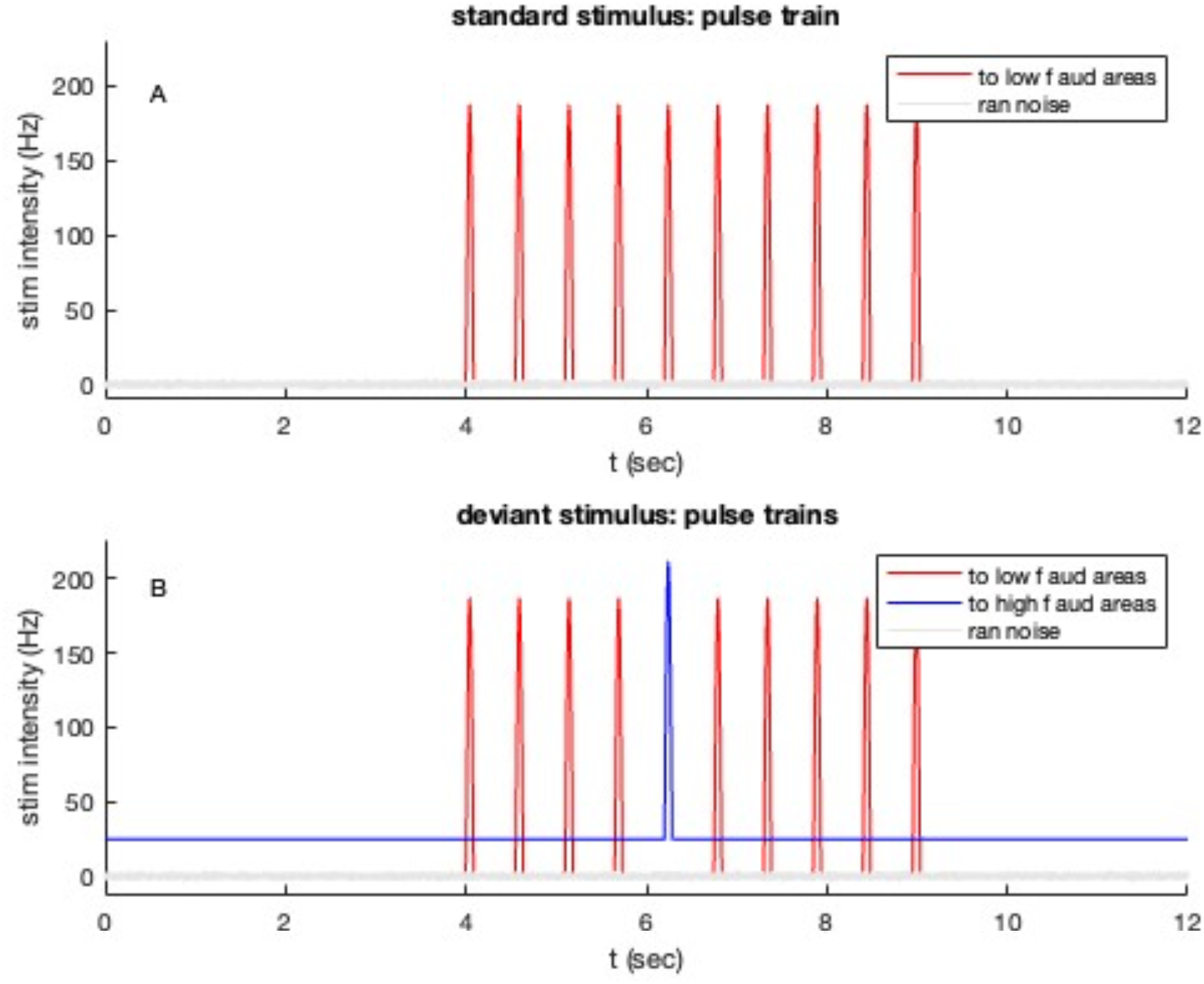
Stimulus function (cf, p(t) in WC/JR models) applied to identified Auditory areas. A) standard stimulus of 10 pulse functions directed to low frequency responsive areas shown in red; B) deviant protocol stimulus, comprising 9 pulse functions (5^th^ pulse absent) to low f areas, and one (5^th^) deviant pulse, shown offset in blue at start time 6.2 s, applied only to high f areas. Details of a single pulse are shown in Supplm. Fig. 1.

Stimuli, illustrated in Figure 1, included 1 (units Hz) standard deviation (sd) Gaussian random noise to all NMs. This protocol is close to that used by Komatsu et. al. (2015) and was more straightforward to implement numerically than the many standards and oddball protocols used experimentally. The shape of the stimulus function is important for numerical stability, and is discussed in the Supplementary Information. A smooth pulse function (cf. Supplm. Fig. 1) of width 70 +14 ms and height 187 Hz produced stable results. Here the deviant pulse has the same width as the standard pulses; comparison with the oddball paradigm is discussed in the Supplementary Information.

For each NM a voltage response is calculated: this follows the standard usage dy = ye – yi, where ye and yi are the outputs of the excitatory and inhibitory neural sub-populations, respectively. The average of contributing dy’s yields the local field potential (LFP) for each cluster. The deviant difference response reported is ΔLFP(t) for each cluster, the difference of LFP(t) in the hi and low sets of stimuli: (9 standard pulses to low f responsive areas, and 5^th^ deviant pulse to high f responsive areas) – (standard stimulus of 10 pulses to low f responsive areas) – cf. Fig. 1. These protocols are referred to as “low” and “hi”, respectively. For each regional cluster the local LFP is the average of dy‘s at participating local areas (i.e. NMs).

## 3. Results

A typical deviant difference response, ΔLFP-Aud(t), to the deviant stimulus protocol (cf. Fig. 1) is shown in Figure 2B, along with the individual standard (to low f areas) and deviant (with high f areas) responses in Fig. 2A. The average deviant response delay over 10 simulation runs was 131.8 ±1.8 (sd) ms for auditory areas and 147.2 ±2.2 ms for pFC. Likely those two timescales relate to signally delays over direct and 2-step pathways, and to phase alignment with target NM oscillations in pFC. A double delay response in frontal areas was also observed experimentally (Komatsu et. al. 2015). All results were statistically robust, using a Wicoxon two sides rank sum test that median potentials under standard and deviant stimuli were significantly different, with p < 4.6 x 10^-62^ for LFP-Aud and p < 0.0034 for LFP-pFC.

**Figure 2.**
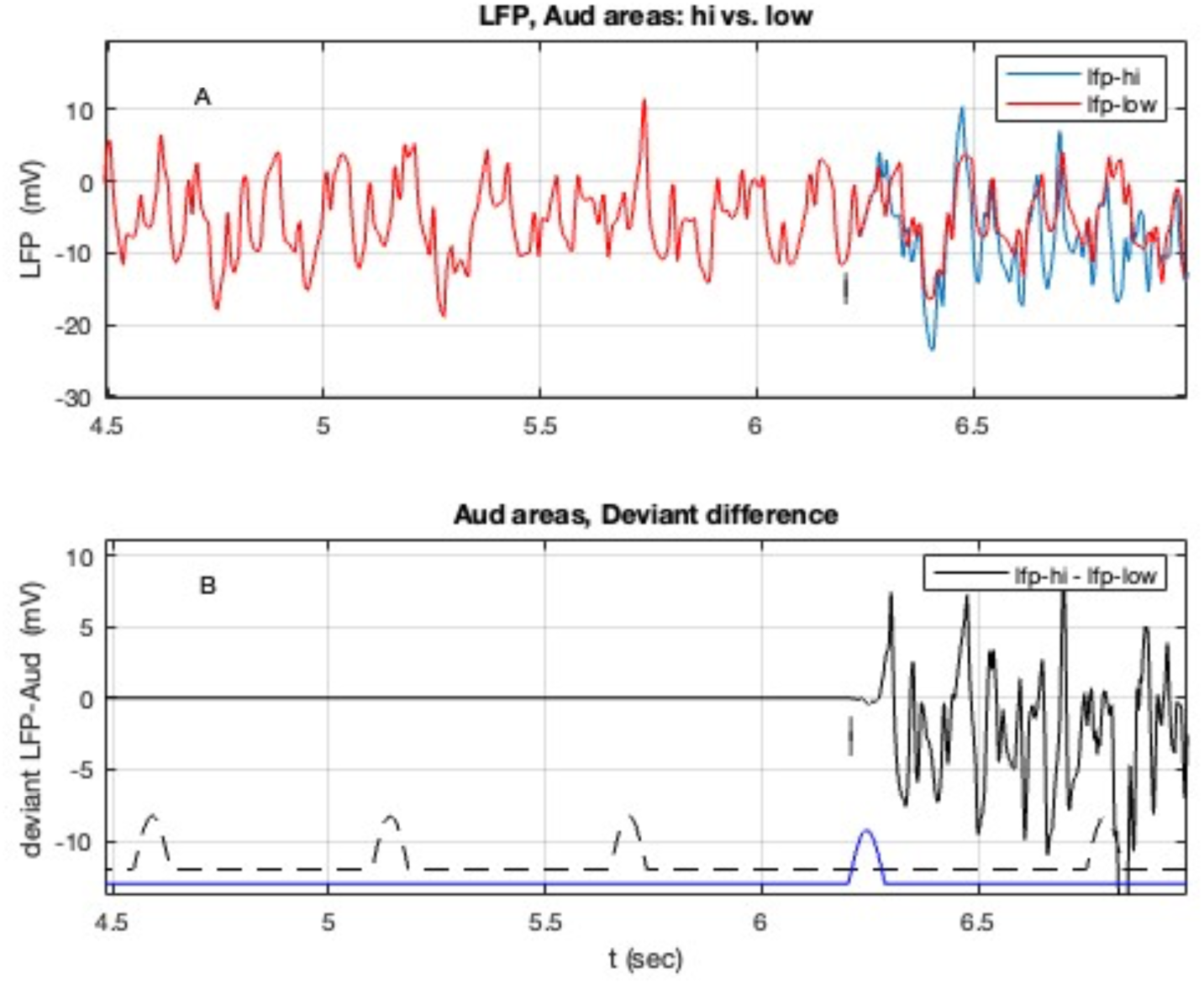
Components of the deviant response in auditory LFP, averaged over all 9 auditory areas in the model. **A**) Top: individual responses to stimuli to high and low frequency responsive areas; standard (low f areas) stimuli are shown in red, with the 5^th^ deviant pulse is shown offset in blue along with its start time (6.2 s) indicated by the vertical mark; **B**) Bottom: Deviant difference response, Δ-LFP, as the difference (high -low) in responses.

These simulation results can be compared to the measured negative ECOG dip at a delay of 46 – 122 ms observed (Komatsu et. al. 2015) near auditory areas; and of 140 – 163, and also 184 – 214 ms at frontal areas; and also to the 100-250 ms range measured experimentally (Obara et. al. 2023). Variability likely arose from the random noise usually applied in the simulations. Local signal delays in each cluster were negligible, but did contribute for pFC given the longer pathways from the auditory areas. Superposition of multiple oscillations in each region (with 9 or 8 areas contributing) also contribute to the final delay (cf. Figs. 7, 12-14).

Similar delay estimates are available for individual neural masses: for Δdy-AuA1 the average deviant response delay was 121.0 ms ± 0.8 (sd); for TPO in two groups: 130.0 ± 1.4 ms and 164.3 ± 1.5 ms – the difference between these two groups is 34 ms, close to the period of gamma band tuning of TPO in the model. For the primary in- and out-hubs in pFC delays are for A32V: 156.5 ± 1.3 ms; and for A10: 140.8 ±2.3 ms.

It is immediately apparent that the oscillations superimpose during the first 4 identical standard pulses, and then start to diverge with a delay from when (6.2 s in Fig. 2A) the 5^th^ deviant pulse is applied and that this gives rise to deviant difference potential response (Fig. 2B), from which the delay time can be measured – as 132 ms in this instance. The negative dip observed experimentally is evident. Note that the published experimental results only plot signal for some 400 ms after the stimulus onset, since they decay thereafter. The general form of voltage responses in Fig. 2A is preserved from one simulation run to the next, with peak to peak amplitudes varying by ∼1-2 mV. The longer time simulated response is more oscillatory and sustained than the apparently smoother experimental observed response. Here multiple cortical columns, that comprise an anatomical area, are taken to be identical NM oscillators with a single frequency. In reality there will be a distribution of characteristics, including oscillatory frequencies, that will yield multiple waveforms that superimpose and tend to smooth out the oscillations shown here (cf. Fig. 7). The corresponding result for pFC is presented in Figure 3.

**Figure 3.**
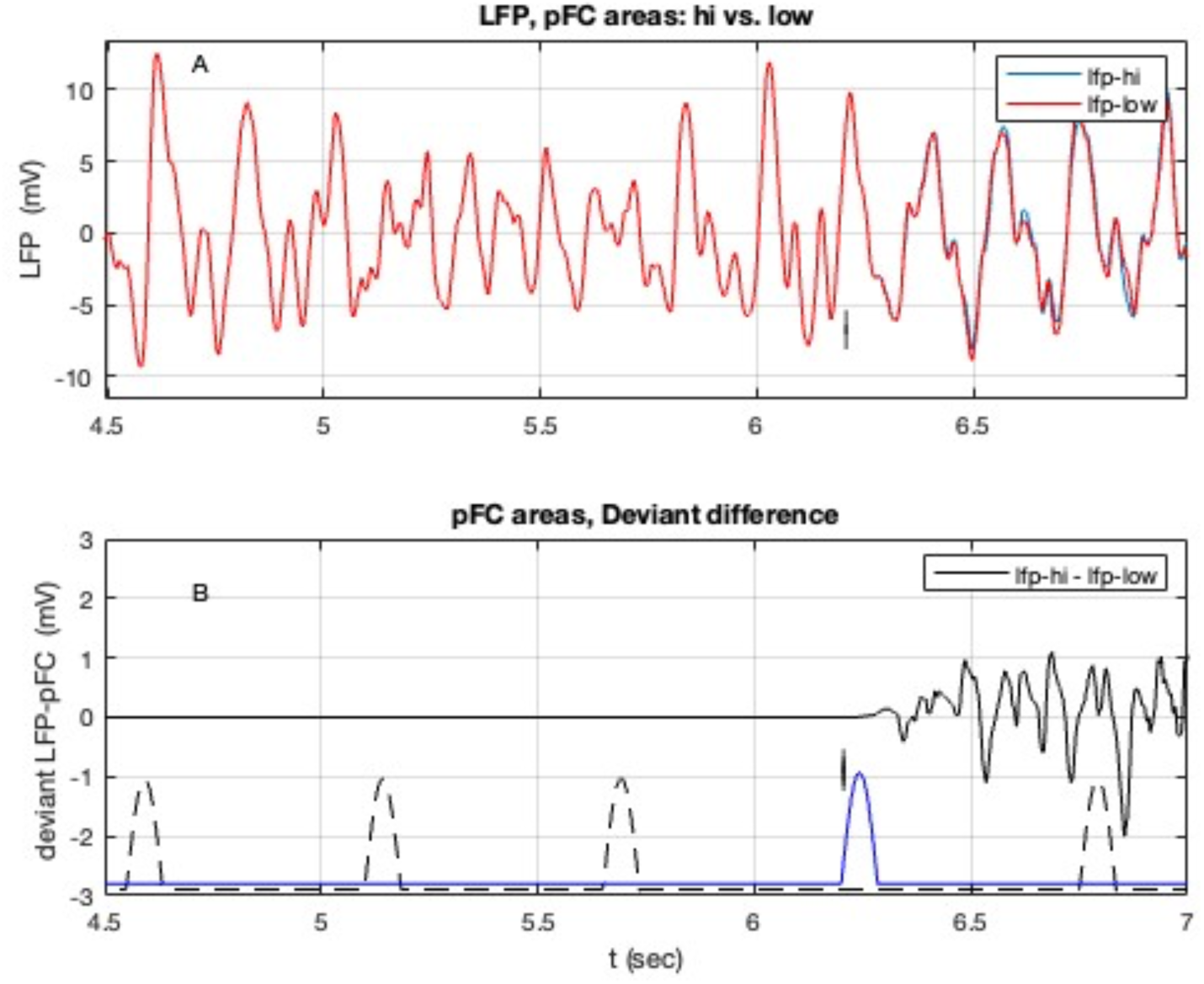
Components of the deviant response in pFC LFP, averaged over all 8 pre frontal areas included in the model. **A**) Top: individual responses to stimuli to standard and deviant stimuli; standard (low f areas) stimuli are shown in red, with the 5^th^ deviant pulse (high f areas) is shown in blue along with its start time (6.2 s) shown by the vertical mark; **B**) Bottom: the difference (high -low) in responses.

Here the deviant response has smaller amplitude with a double dip and is harder to discern, likely due to the multiple signalling pathways from the auditory to the pFC cluster. The first dip in deviant potential is to -0.41 mV at 145 ms after the deviant stimulus pulse onset (vertical marks in Fig. 3). The extra 15 ms delay is consistent with auditory – pFC path lengths.

### 3.1 Responses in Auditory areas

Individual components of LFP, the dy’s for each anatomical area are also available from the simulations. These are the neural mass voltage responses for component areas. TPO (in the temporal lobes) has the strongest links forward to pFC, into A32V and appears to be a key output node from the auditory cluster. TPO’s individual deviant difference response, Δdy, is shown in Figure 4, along with the individual low and high responses: the delay in deviant difference response is 129 ms. That is 9 ms longer than the delay in AuA1 (cf. Fig. 6). Comparing the responses to standard (“low”) and deviant (“hi”) stimuli illuminates the origin on the deviant difference delay. Initially its due to a phase shift in the voltage waveform. Due to the differing frequencies present the phase (mis-)alignment drifts and, additionally, differences in amplitude become apparent. At longer times (cf. Supplementary Figure 2) the 6^th^ standard pulse (at 6.75 s) takes effect and induces an altered pattern with a secondary deviant difference delay (125 ms, vs. 129 ms for the deviant stimulus).

**Figure 4.**
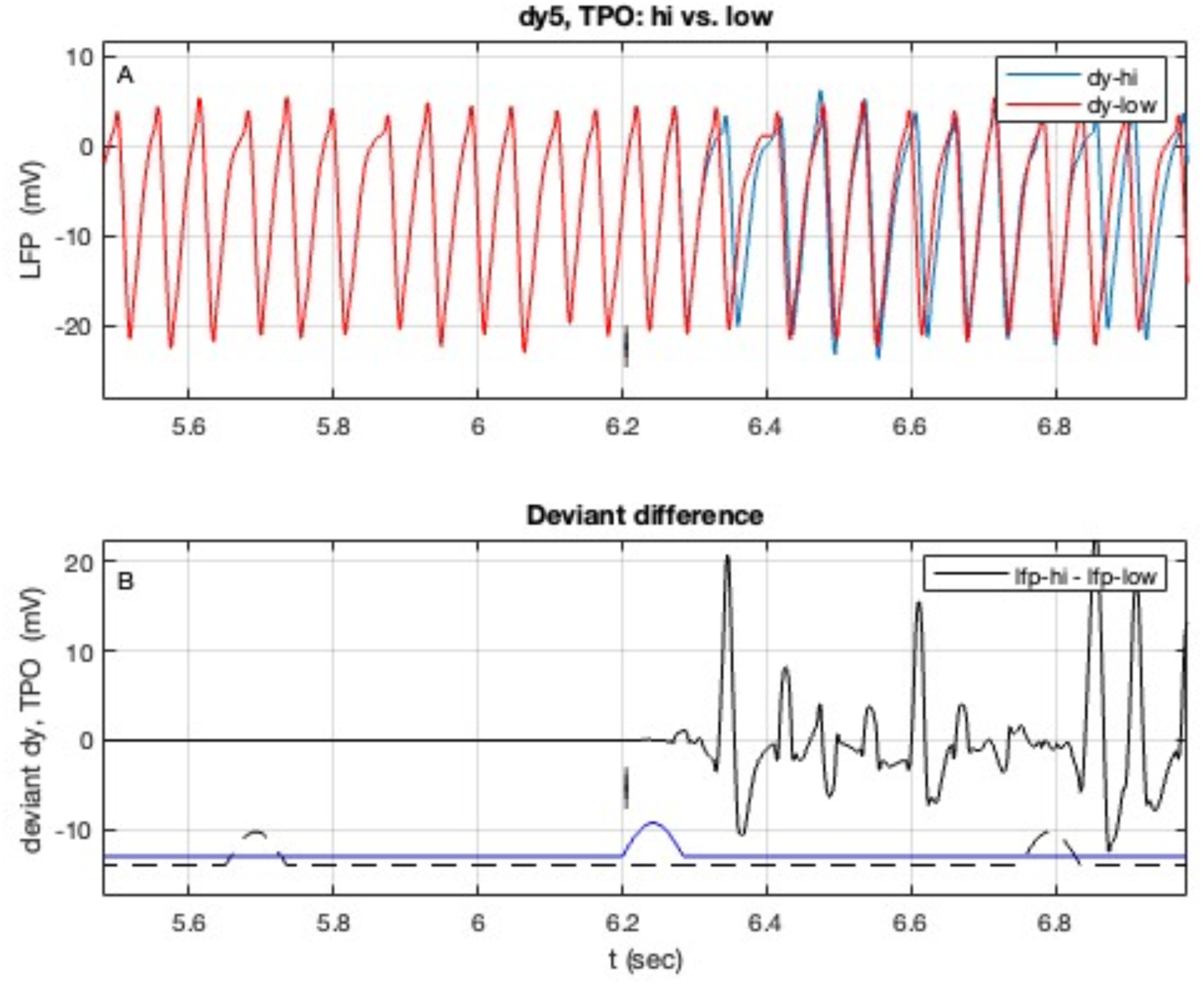
Deviant response in area TPO. **A**) individual responses to stimuli to standard (“low”) and deviant (“hi”) stimuli; **B**) the deviant, ie. the difference in responses in A. Details as in Fig. 2.

Area AuRPB has been identified as a critical link in the NMM response (Obara et. al. 2023 **check**). Its response is shown in Figure 5, along with the individual low and high responses. The delay, in this instance, is 126 ms. The longer time course of Δdy-AuRPB is presented in Supplm. Fig. 3 and shows an interesting symmetric bursting that decays for 4 s after the stimulus pulses ends. The overlay of dy-hi(t) and dy-low(t) in Fig. 5A highlights the origin of the dip in Δdy (cf. Fig. 5B) as being the upward displacement of the voltage dy(t) due to the deviant stimulus. This is also evident in the other results. AuRPB has quite strong links back to the core: RPB - RT has weight 2.1k (see Pailthorpe 2023a for detailed discussion of link weights) and RPB - CL 1.0k, consistent with the strong feedback noted experimentally (Obara et. al. 2023).

**Figure 5.**
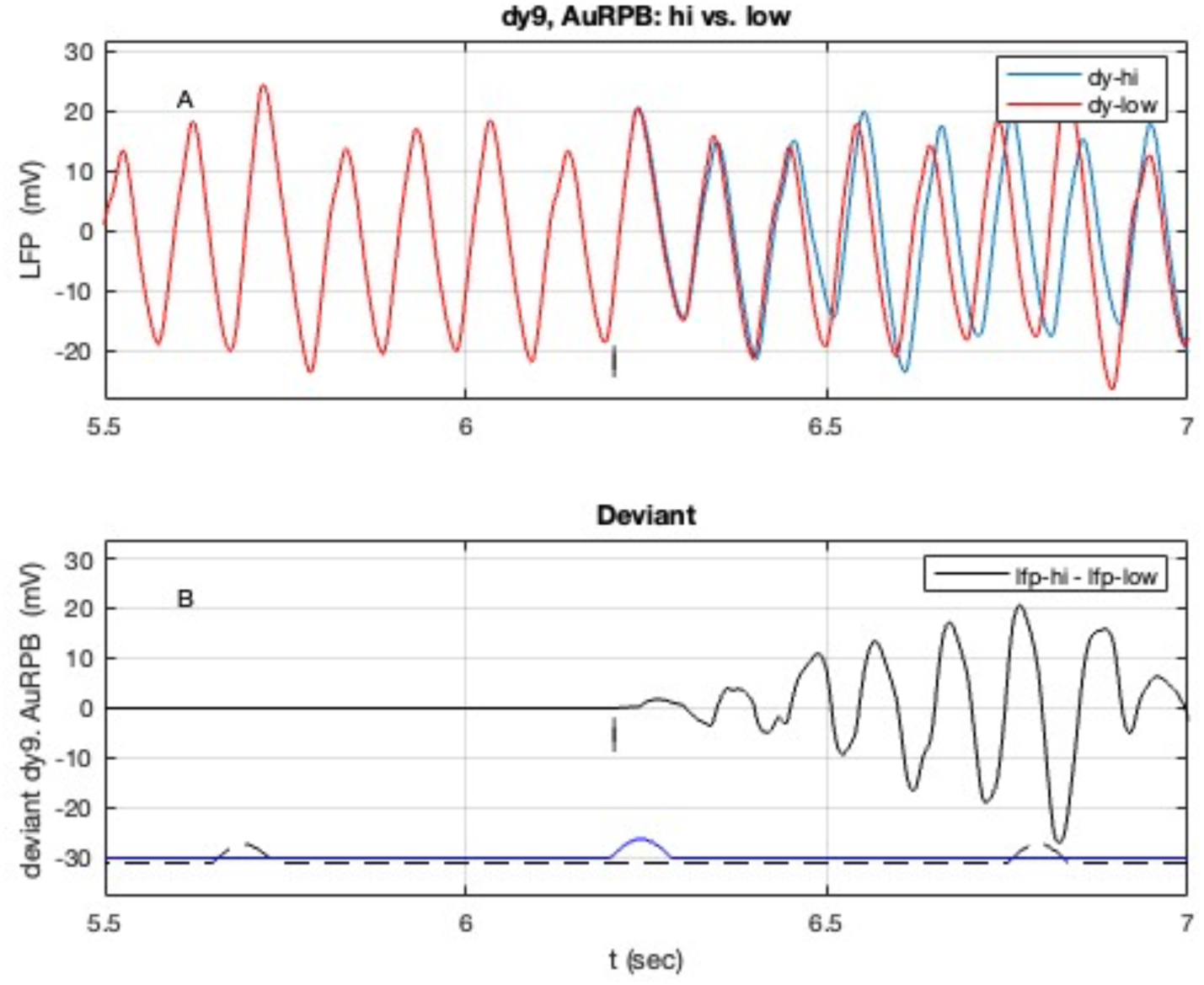
Deviant response in auditory area TPO. **A**) individual responses to standard (“low”) and deviant (“hi”) stimuli; **B**) the difference in responses. Details as in Fig. 2.

The primary auditory cortex area, AuA1, is a key initial target and intermediary in the auditory response pathway. It is evident here that AuA1 only oscillates when an external stimulus is applied. It has a weaker response to “low f” stimuli than to “high f” (cf. Methods). Its deviant response starts oscillating following the 5^th^ deviant pulse, as shown in Figure 6 along with the individual low and high stimulus responses. The delay is 120 ms. AuA1 has numerous strong out links, the strongest being locally to RT (weight 2.8k), and RPB (1.8k) and to TPO (1.8k). These links mentioned are amongst the strongest links in the auditory cluster and highlights the key roles of those nodes. TPO was studied herein since it receives strong (weights > 1k) inputs from A1, CL, CL and CPB, and a medium strength (0.4k) input from RPB. It also has medium weight links forward to pFC (weights 0.6k, 0.5k, 0.3k).

**Figure 6.**
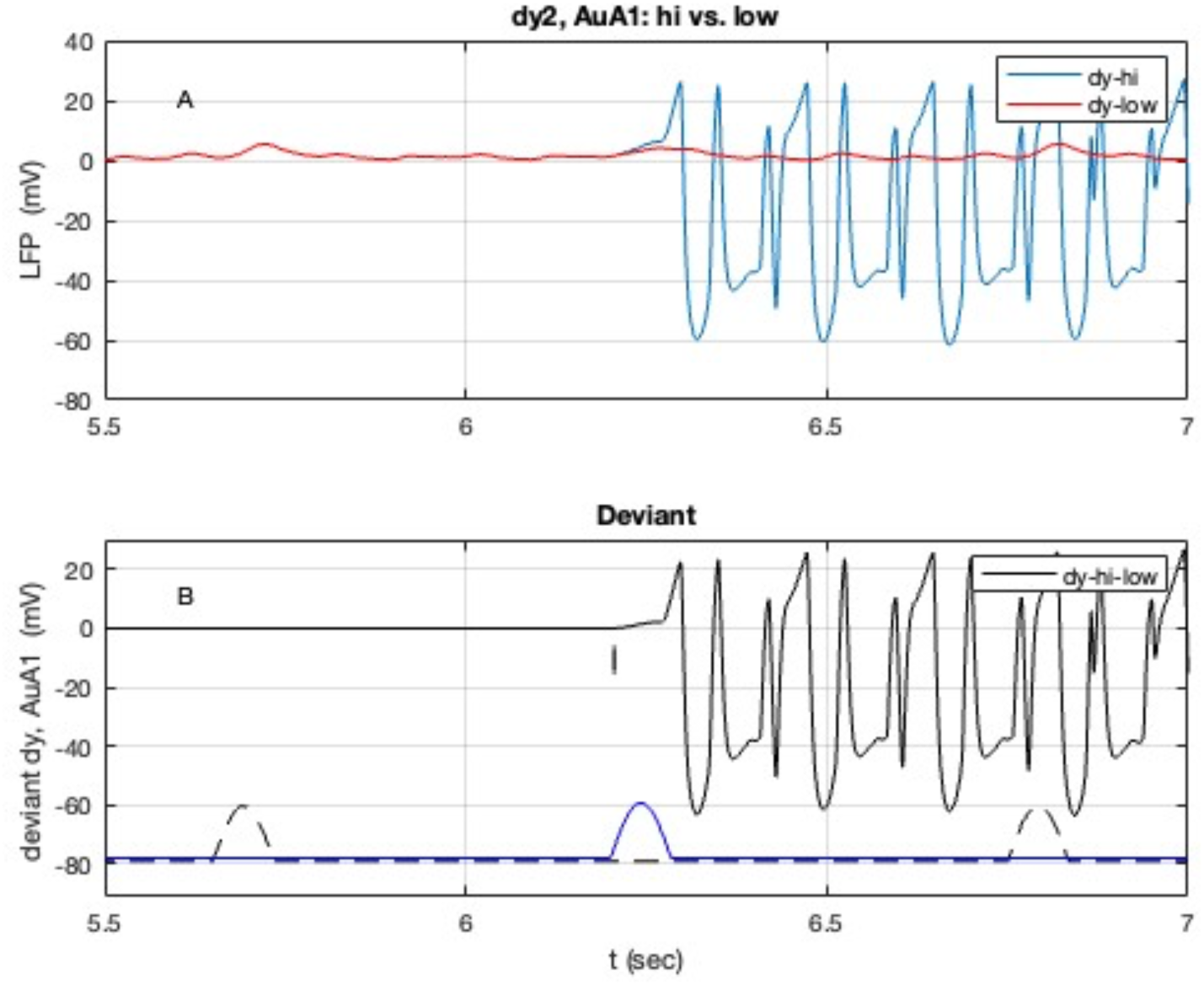
Components of the deviant response in auditory area AuA1. **A**) individual responses to stimuli to high (deviant) and low (standard) frequency responsive areas; **B**) the difference in responses. Details as in Fig. 2.

The auditory cluster LFP (Fig. 2) is a complex waveform, while the individual area’s LPFs are simpler oscillations (some simply sin waves), reflecting the characteristic frequency of each area (Pailthorpe 2024b). The LFP is a composite (here computed as a simple average) of the constituent dy’s, shown below in Figure 7 - they superimpose, adding or subtracting to constructively or destructively interfere, and so produce the composite LFP shown in Fig. 2.

**Figure 7.**
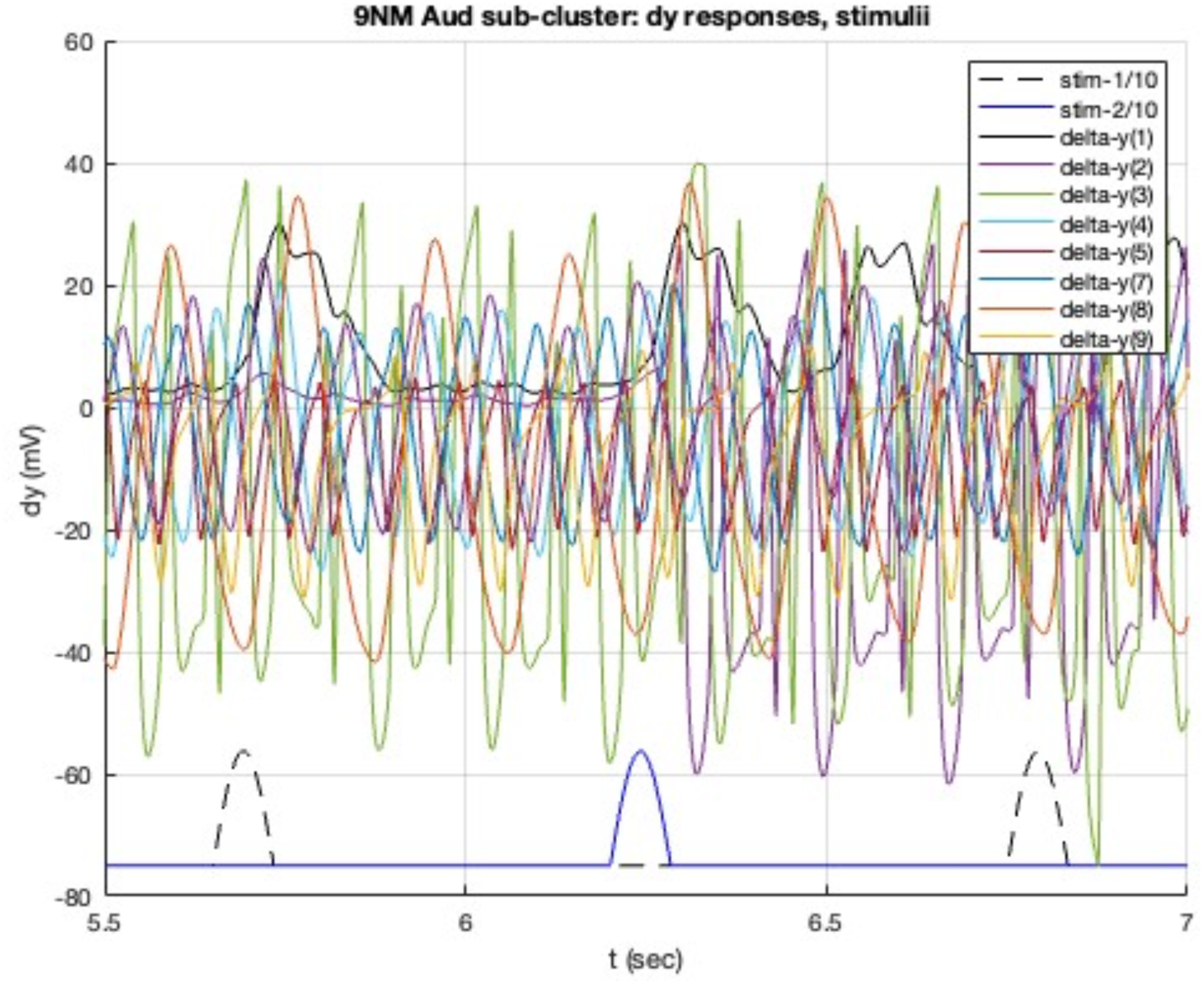
Component dy’s of the auditory LFP (Fig. 2) for the nine neural masses used to model the auditory cortex, under a deviant stimulus. The 4^th^ and 6^th^ (in a stream of 10 pulses) low f stimulus pulses are shown (dashed black line) below, along with the 5^th^ deviant pulse (solid blue), as in Fig. 2.

This shows that most oscillations of the NM potential are simple sinusoids, with some being slightly more complex mixtures. Their superposition yields the more complex waveform of the Auditory LFP shown in Fig. 2. It is evident that some nodes respond promptly to the stimulus pulses. Note that AuA1 (dy-2, purple in Fig. 7) exhibits a relatively high amplitude response to the 5^th^ deviant pulse. The delay in LFP, which is a superposition of the oscillations in Fig. 7, emerges from the constructive and destructive interference of these individual oscillations. The observed frequencies in the simulations are listed in Table 1, spanning beta to gamma bands: 5.5; 9.7, 9.9; 15.2, 16, 18.3; 32.4 Hz – corresponding to periods of 182; 103, 101; 66, 63, 55; 31 ms. These oscillation periods T are credibly in the range of observed delays. Zooming in on Fig. 7 and comparing with the resultant LFP-Aud, as displayed in Figure 8 shows the response of individual areas. The delayed dips in LFP-Aud after the pulse at 6.2 s (Fig. 8A), along with the corresponding large dip in dy-A1 (Fig. 7 and 8B), are evident. This highlights how AuA1 has differential sensitivity to low and high f stimuli. Essentially AuA1 turns on with the deviant stimulus.

**Figure 8.**
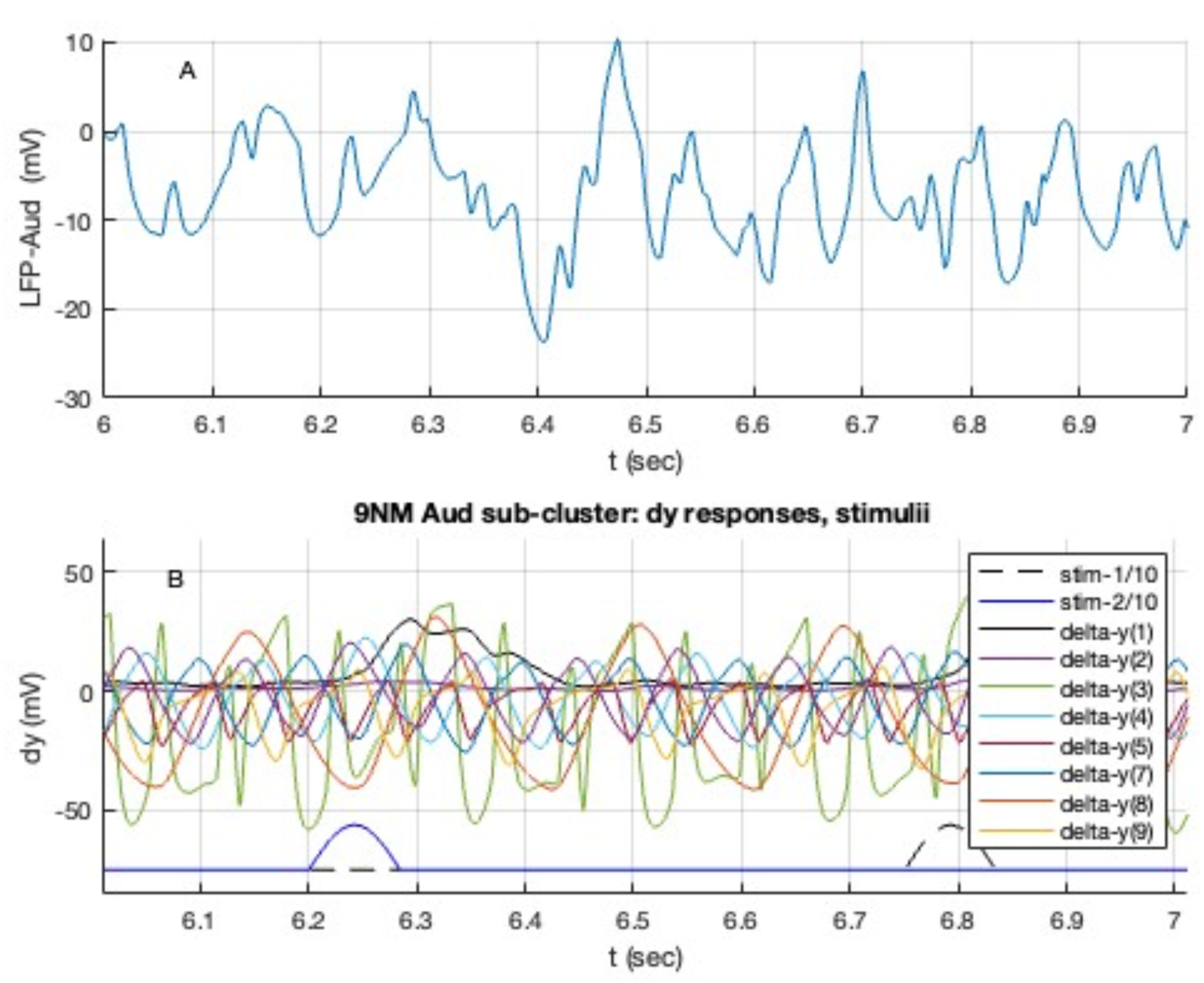
**A**) auditory LFP (cf. Fig. 2) and **B**) its component dy’s (cf. Fig. 7) for the nine neural masses used to model the auditory cortex under a deviant stimulus (cf. Fig. 1). The 6^th^ (in a stream of 10 pulses) low f stimulus pulse is shown (dashed black line) below, along with the 5^th^ deviant pulse (solid blue), as in Fig. 2. Deviant stimulus onset at 6.2 s.

The change in signal phase can be studied using the analytic signal, calculated via Matlab’s Hilbert transform with *designfilt* and *filtfilt* functions with a zero-phase bandpass filter in 0- 500 Hz. For the deviant signal from the NM representing AuA1 the time dependent phase is shown in Figure 9. Standard stimulus pulses increase the phase from ∼ 0° progressively up to near 90°. The 5^th^ deviant pulse resets the phase to near 0°, with a larger variance that decays progressively to a stable, but non-zero, range. The strong out-link from AuA1 are to: AuRT locally in the core, then to AuCM and TPO; to AuRPB and AuCL.

**Figure 9.**
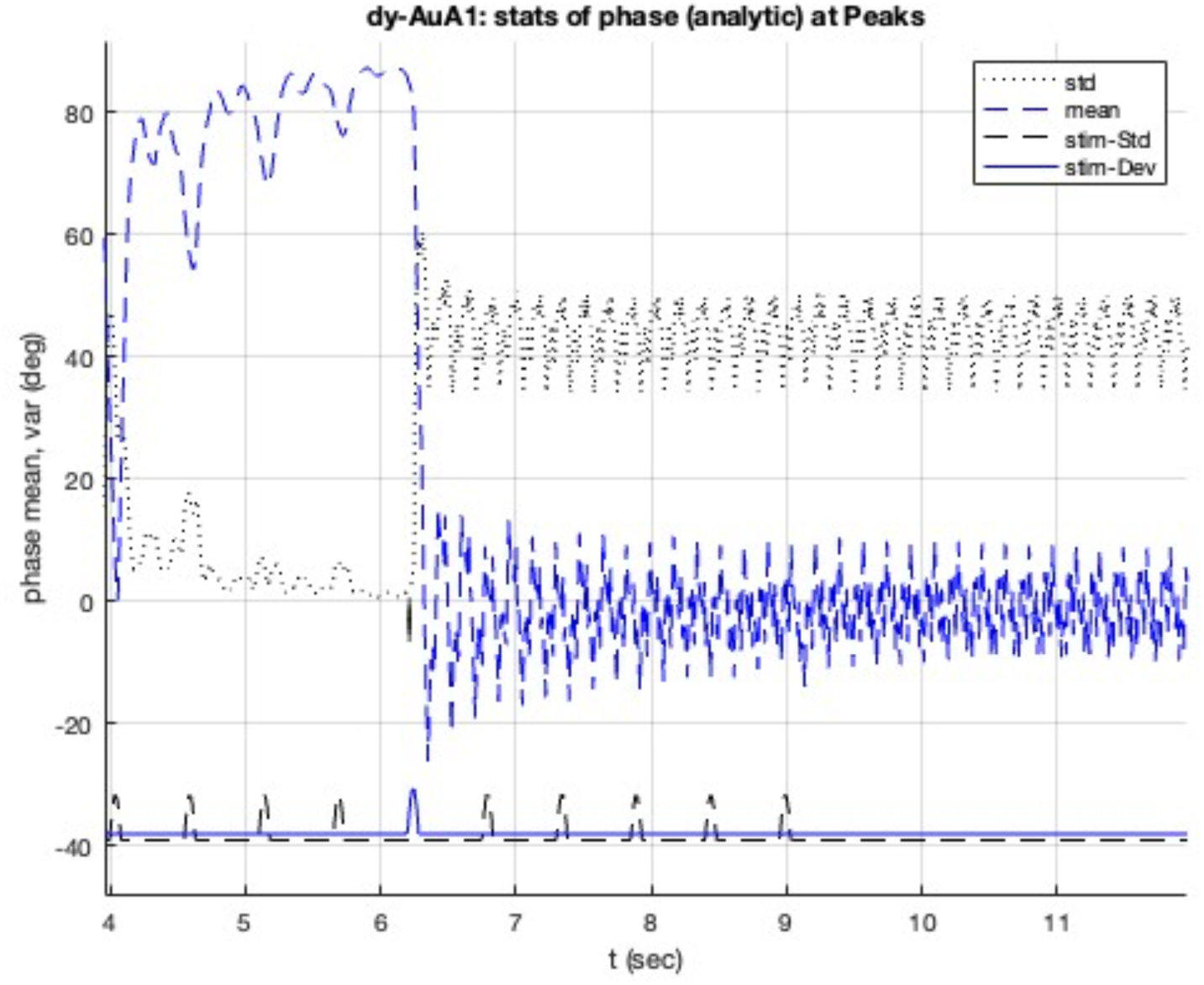
Moving average of the analytic phase of deviant response in auditory area AuA1 (cf. Fig 6A); variance is also plotted. Stimulus pulse plotted at bottom, as in Fig 2.

### 3.2 Responses in pFC

One presumed target of auditory responses is pFC, which is explicitly modelled here using the two interlinked clusters. The LFP for the 8 nodes comprising pFC is shown in Fig. 3. The key in-hub is A32V (P 2023a, 2023b), which also hosts the strongest incoming links from TPO in the auditory cluster (cf. Fig. 4). In pFC areas A32, A9 show no evident response, having little if any direct inputs from auditory areas. A8aD and A8b have relatively weak responses but appear to be required participants given that they receive other sensory inputs and have links to pre motor areas. Those 4 areas receive inputs via the waypoints discussed above. Key responses are in the more strongly linked hub nodes, A32V, A11, A10, along with A46D. The in hubs, A32V and A11, show sustained bursting patterns, as displayed in Figures 10 and 11. For A32V, tuned in the theta band (cf. Table 1), the deviant induced oscillations show multiple spectral peaks, dominated by 5.19 and 10.4 Hz; spectral power is 1% delta, 82% theta, 13% alpha, 4% beta and 0.4% gamma. The long time deviant difference response in A32V (Fig. 10) is small oscillations after a delay of 158 ms that continue until interrupted by the 6^th^ to 8^th^ standard spikes to low f areas. Subsequent negative spikes are larger and continue up to 9.5 s, some 0.41 s after the end of the stimuli (and 0.54 s after onset of 10^th^ std pulse). There follows a relatively quiet period up to 11.2 s (at 2.1 s after end of Pulse #10), followed by a sustained burst of positive spiking. For A11, tuned in the beta band, the oscillations are more symmetric, with sustained ringing, and show two spectral peaks at 15.56 and 32.8 Hz, with spectral power 90% in the beta band and 8% gamma.

**Figure 10.**
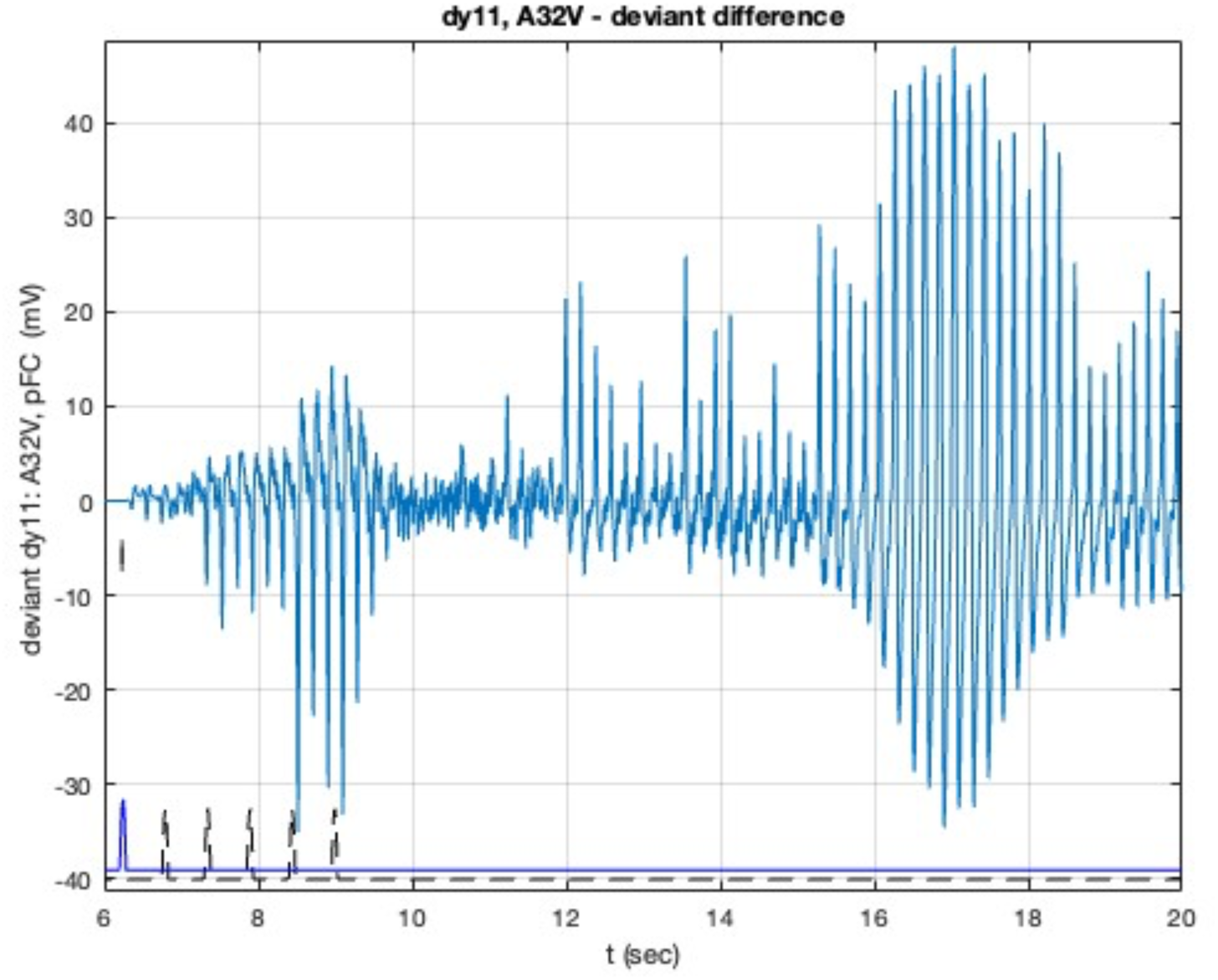
The long term deviant difference response in pFC area A32V, the primary in-hub in pFC. Standard (to low f areas) stimuli are shown in black, with the 5^th^ deviant pulse shown offset in blue along with its start time, 6.2 s, indicated by the vertical mark.

Interestingly pFC area A10 was identified as a key out-hub in the marmoset network connectivity data, and so was examined in my earlier study of a pFC model (P 2023a, 2023b). Its strong out links are to: nearby pFC (A47L, A45), then motor areas, then feedback to auditory areas (TPO, AuCPB, AuML, AuCM), to cingulate areas (A23a, b), to multiple PPC areas, and to TE3 in lateral temporal cortex. Its initial deviant response was -0.8 mV at a delay of 141ms (past the onset of the 5^th^ deviant stimulus pulse). Subsequently it exhibits a train of 9 gamma bursts (initial period 31 ms, f = 32.3 Hz) in response to the deviant stimulus, as shown in Figure 12. The decay of that burst is interrupted by the next (6^th^) standard stimulus pulse to low frequency sensitive auditory areas that induce a new, stronger and longer burst at the same frequency. That burst decays until again regenerated by the next stimulus pulse. With subsequent pulses, and increasing energy input to the NM oscillators, the oscillations grow larger, and then decay some 1.1 s after the onset of the 10^th^ pulse at 8.95 s. The general features of this bursting are preserved, with its details varying from one simulation run to the next, likely due to the bursting being sensitive to the random noise input to the multiple NMs – cf. discussion around Supplm. Fig. 11.

**Figure 11.**
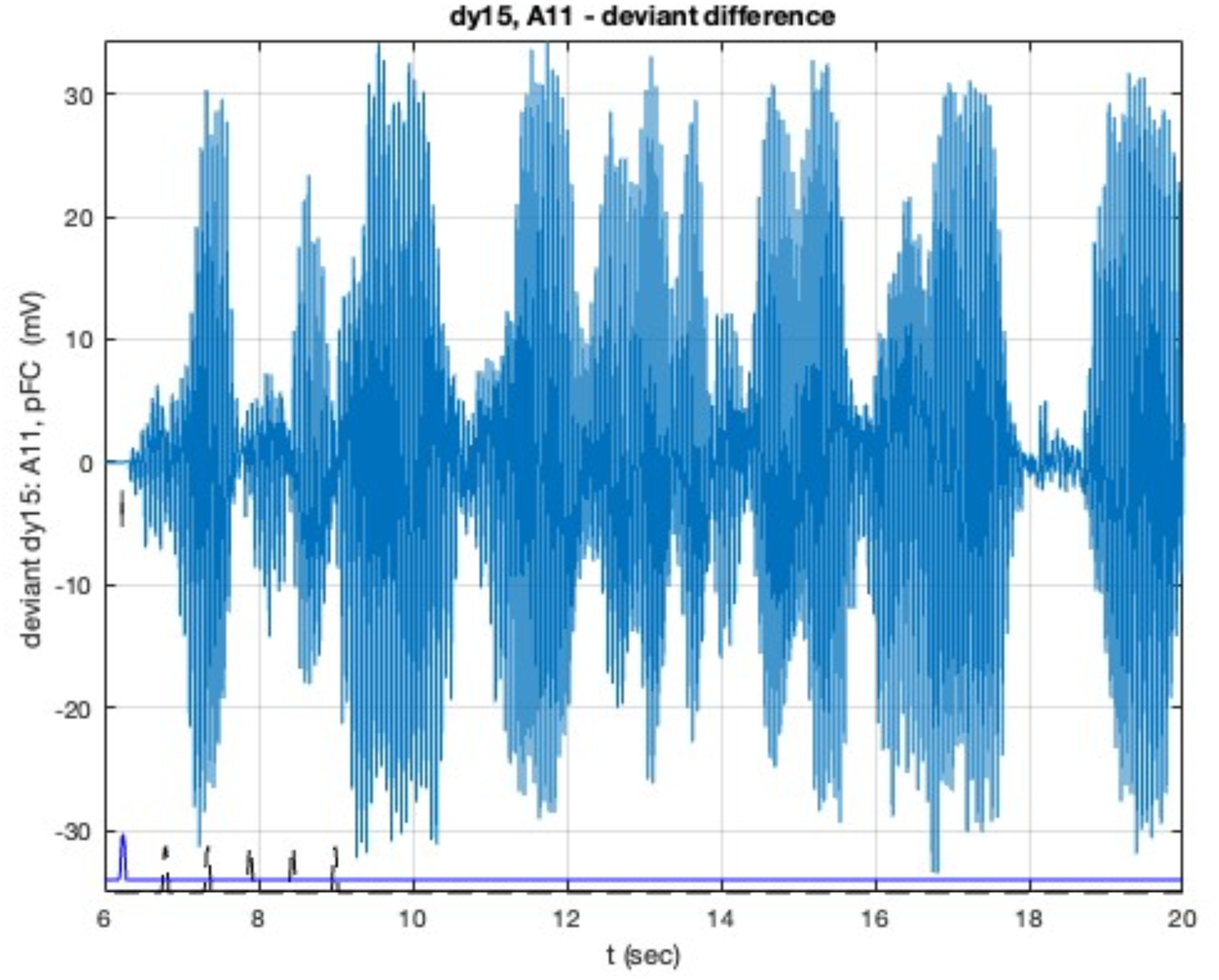
The long term deviant difference response in pFC area A11, the secondary in-hub in pFC. Details as in Fig. 10.

**Figure 12.**
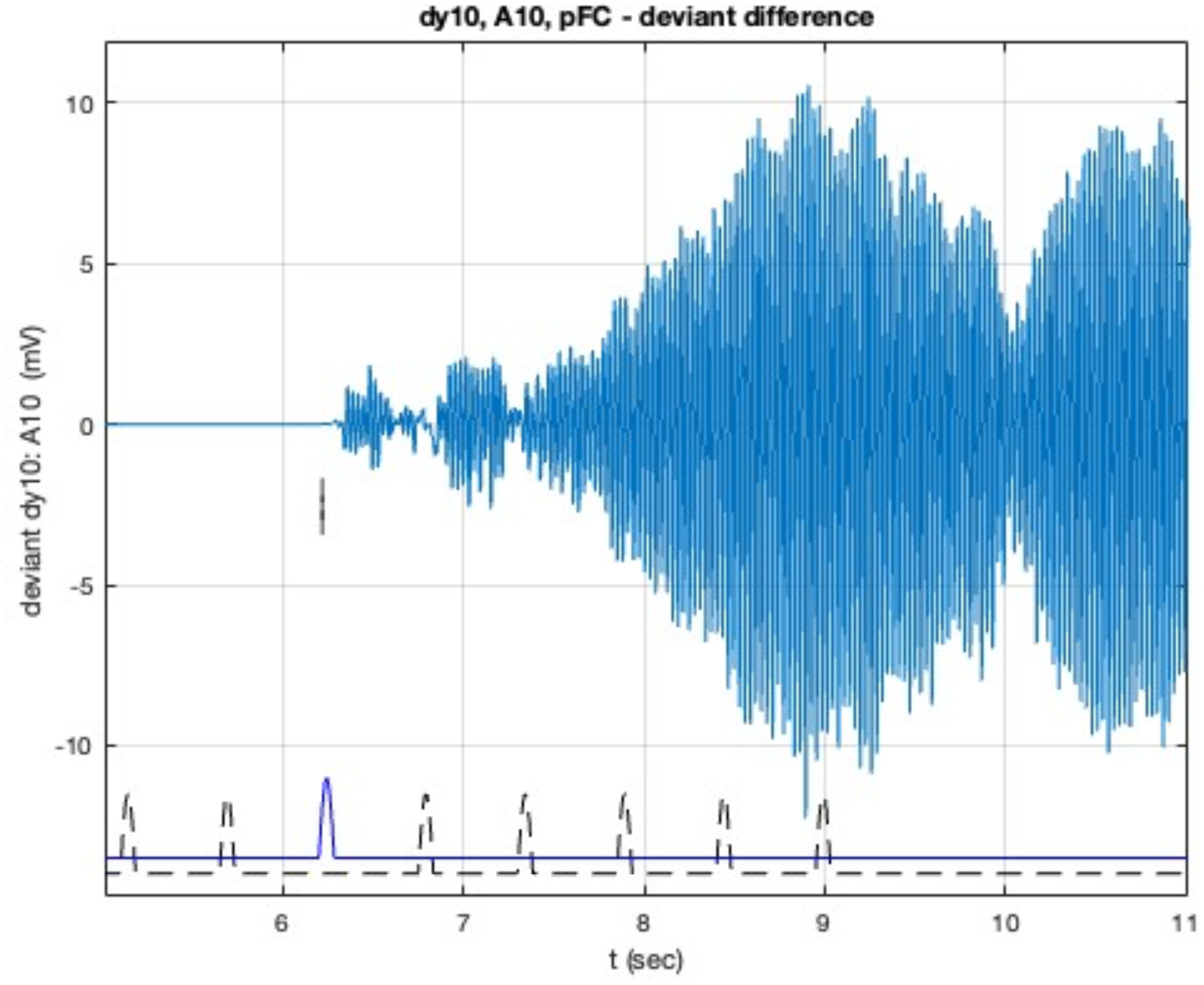
An example of the deviant difference Δdy response in the pFC out-hub, area A10, calculated as the difference (high -low) in responses to stimulus pulses to high and low frequency responsive areas. Standard (low f areas) stimulus pulses are shown (dashed black line) at the bottom, with the 5^th^ deviant pulse (offset, solid blue) with its start time (6.2s) indicated by the vertical mark. Other details as in Fig. 2.

The fft Power Spectral Density of this response (over 4-20 s) has a single peak at 32.8 Hz (period 30.5 ms), with 99% of spectral power in the gamma band and 0.8% of in the beta band.

### 3.3 Details of the Auditory Response

The response of the 9 nodes were plotted together in Figs. 7 and 8. Examining now the responses in each of core, belt, parabelt and temporal regions sheds more light on how individual waveforms contribute to the deviant difference delay. The response to the deviant stimulus (cf. Fig. 1) of NMs in the core are shown in Figure 13.

**Figure 13.**
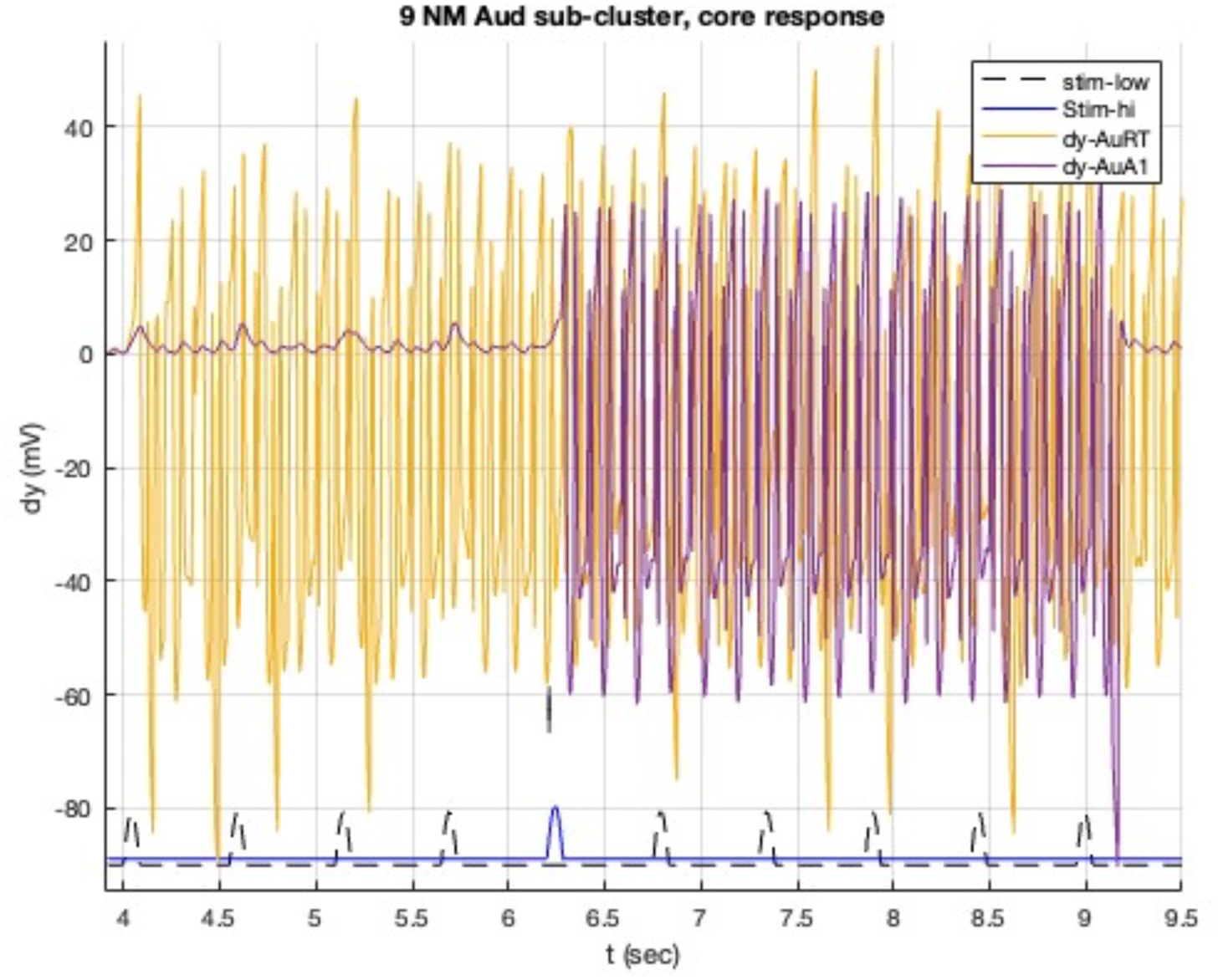
Individual voltage responses of neural masses in the core of the auditory sub-cluster, under a deviant stimulus. The standard stream of 10 stimulus pulses to low f sensitive areas are shown (dashed black line) below, along with the 5^th^ deviant pulse (solid blue, offset) to high f sensitive areas, as in Fig. 2.

None of the auditory NMs oscillate prior to the external stimuli. AuRT, tuned in beta/gamma bands, responds quickly, after 63 ms, to the standard stimulus pulses (to the low f responsive areas) and exhibits sporadic fast spikes; it responds less to the deviant pulse. AuA1 responds weakly to the standard stimuli, and strongly only after the deviant stimulus (5^th^ pulse) to the high f responsive areas, and then shuts down.

Responses in the belt areas are shown in Figure 14. The time axis is expanded to discern individual waveforms. AuAL responds preferentially to standard stimuli. AuML shows a stronger response to the deviant, vs. standard, stimulus pulses. AuCL has a distinct response to the deviant (5^th^) pulse. Note that AuCM was not subject to the stimuli absent information on its responsiveness.

**Figure 14.**
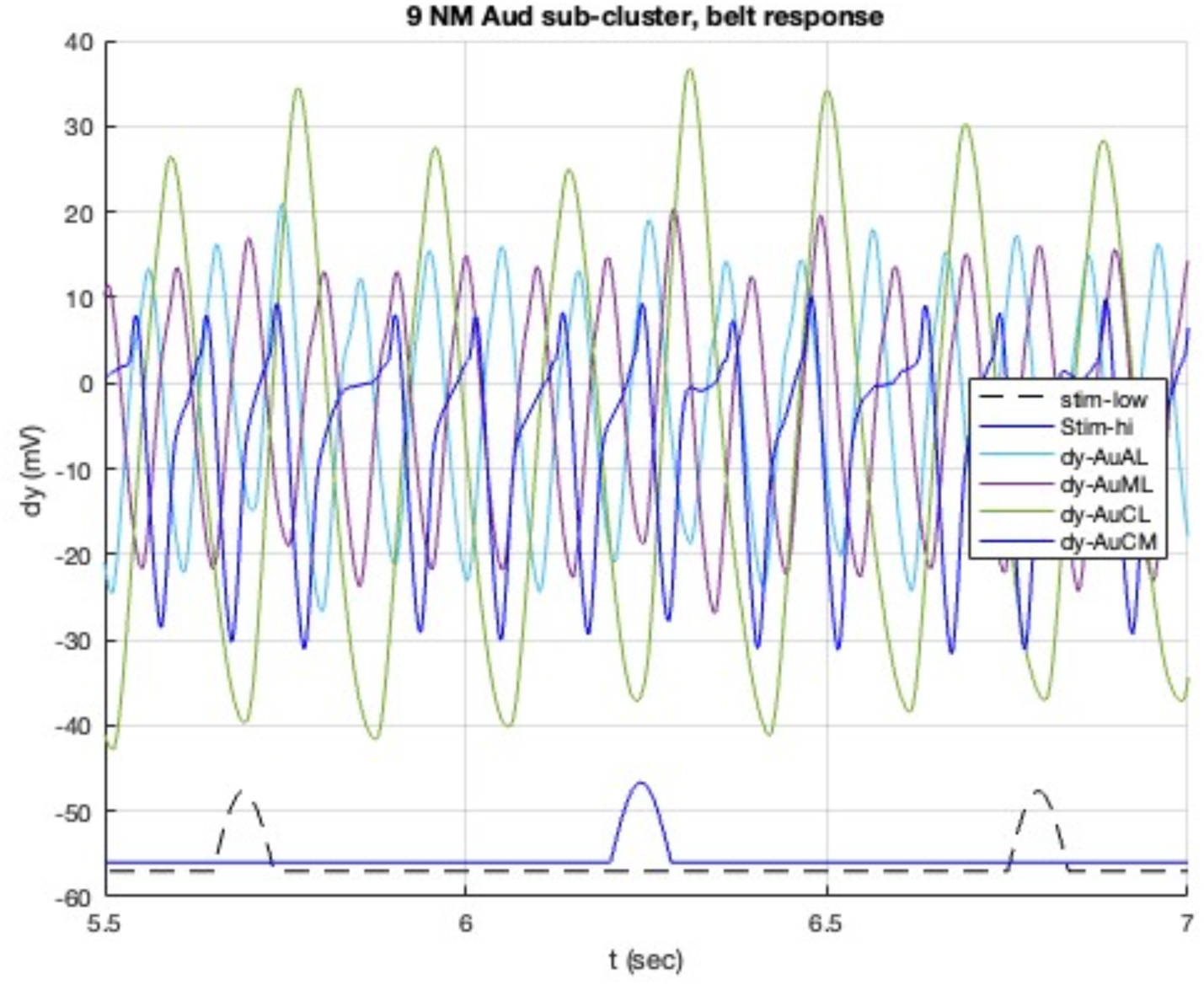
Individual voltage responses of neural masses in the belt region of the auditory sub-cluster, under a deviant stimulus. The standard stream of 9 stimulus pulses to low f sensitive areas are shown (dashed black line) below, along with the 5^th^ deviant pulse (offset, solid blue) to high f sensitive areas, as in Fig. 2.

Responses in the parabelt areas are shown in Figure 15. Both AuRPB and AuCPB have comparable responses to low and high f stimuli. Yet AuCPB, modelled by a 3- and 5-sub- population NM, has a distinct response to the 5^th^ deviant pulse which induces multiple bursts. TPO, while a temporal area, appears naturally in the auditory cluster due to its close network integration, so its response is included here. TPO and AuCPB have the strongest onward links to the pFC cluster.

**Figure 15.**
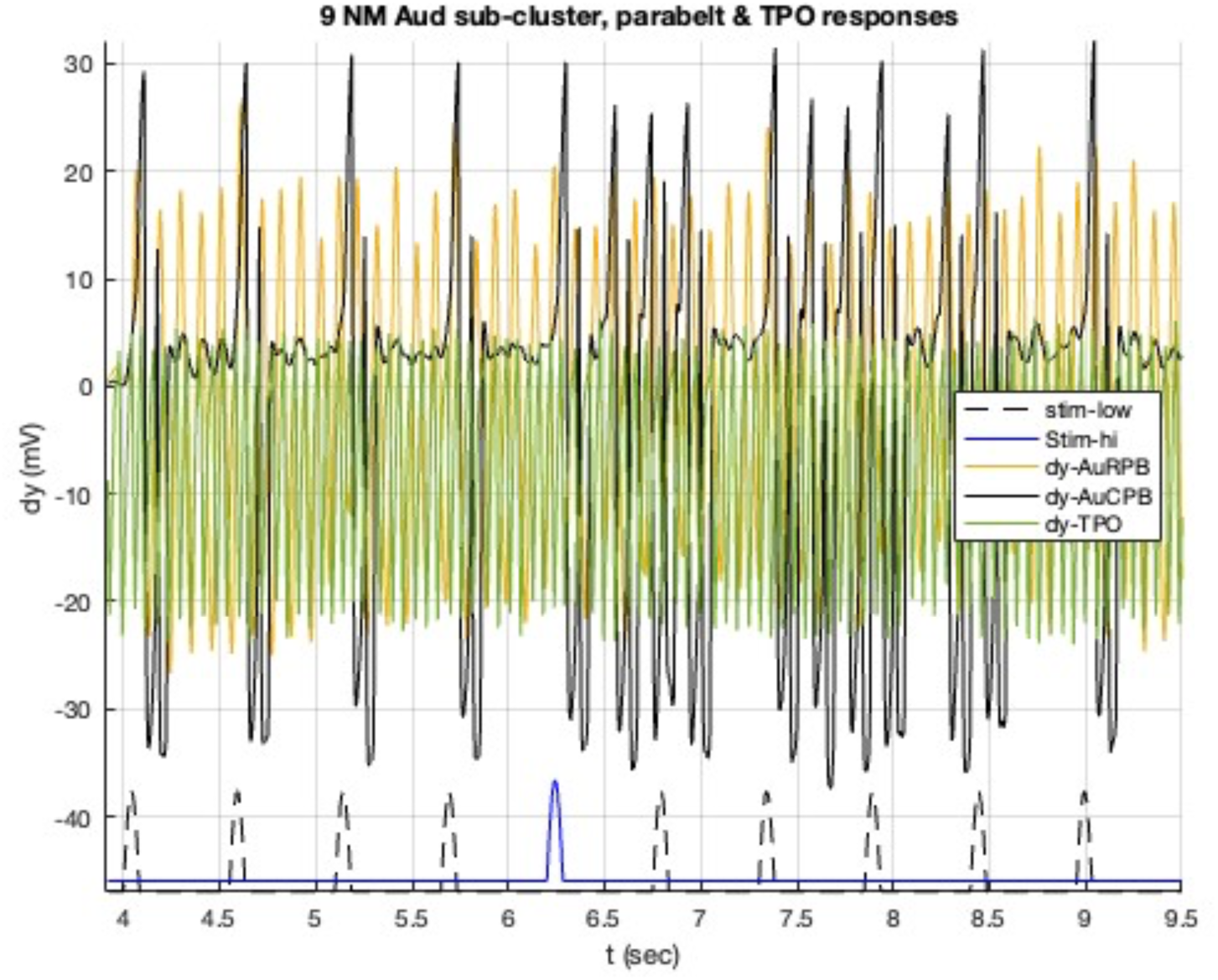
Individual voltage responses of neural masses in the parabelt region of the auditory sub-cluster along with TPO, under a deviant stimulus. The standard stream of 9 stimulus pulses to low f sensitive areas are shown (dashed black line) below, along with the 5^th^ deviant pulse (offset, solid blue) to high f sensitive areas, as in Fig. 2.

AuRPB oscillates throughout and responds with a single enhanced spike to the initial standard pulses, but not to the deviant pulse. AuCPB has a burst response to standard stimulus pulses, and then shows multiple bursts in response to the deviant pulse – these decay eventually. TPO, the key outward link to pFC, receives multiple strong inputs from the other auditory areas. It responds with an enhanced double oscillation after the deviant pulse.

### 3.4 Role of AuRPB

Area AuRPB has been identified as a key participant in the deviant NMM response [Obara et. al. 2023], so that was tested by inactivating NM node #9, representing AuRPB, by setting the WC/JR feed-forward and -back coefficients A, B and C to zero, resulting in zero voltage responses in the NM representing AuRPB. As expected, this changes some of the responses discussed above, with some unfamiliar, or ambiguous, oscillatory patterns emerging - a clear NMM response is absent. ΔLFP-Aud now exhibits an altered, faster response, with rapid dips at delays of 116 ms, compared to the average delay of 132 ms in the regular deviant protocol (cf. Fig. 2). The primary auditory area, AuA1 is again activated by the 5^th^ deviant pulse with a faster response, comprising a similar voltage dip at 108 ms after the 5^th^ deviant stimulus onset – earlier than in the usual deviant protocol (cf. 120 ms delay in Fig. 6) - repeating at 18.5 Hz. The response in AuCPB is relatively unchanged by inactivation of AuRPB, consistent with observation. It also is activated by the 5^th^ deviant pulse and exhibits a double spiking pattern with each subsequent pulse, and then is quiescent after stimuli cease.

A noticeable difference is in the deviant difference response of TPO, the key link from the auditory cluster on to pFC areas, as shown in Supplm Figure 7. The response is reduced in amplitude and follows a different time course: there is a small (1/3 amplitude) dip at a delay of 45 ms after deviant onset, vs. 131 ms in the regular case (cf. Fig. 4). A noticeable difference is in phase alignment (Supplm. Fig. 7A), which compares the deviant and standard responses and is illustrative of underlying mechanisms that generate the deviant difference in the present model. Here a 11 ms phase lag (deviant -standard) appears 5 oscillations after deviant onset, vs. the deviant dy(t) leading the standard dy(t) by 3 ms after 8 cycles following the deviant pulse in the regular protocol, as shown in Fig. 4. There appears to be a reduced response to the stimulus pulses.

With TPO being the key onward link to pFC the new output then changes the response in pFC. The deviant difference ΔLFP-pFC does not show a recognisable dip until 173 ms after the deviant stimulus onset, and the waveform appears to be more noisy – this likely just marks the onset of local oscillations in the in-hubs. Δdy-A32V, the main pFC in-hub, shows a slight dip at 192 ms after deviant onset, but has on small responses to the stimulus pulses Similarly Δdy-A11, a secondary pFC in-hub, exhibits a growing oscillation 323 ms after deviant onset. Again, this could be related to local oscillations.

The other parabelt area modelled herein, AuCPB (NM #1), does have a stronger link back to AuRT (NM#3), of weight 2084, but weaker feedback to AuA1, of weight 48, thus limiting its influence. Inactivation of AuCPB, NM#1, was also investigated by setting its parameters A, B, C, G = 0, resulting in zero voltage output for that NM. Response in the core (AuA1, AuRT) was relatively unchanged. Overall ΔLFP-Aud has a slightly altered waveform, now absent the contribution from AuCPB, with a similar amplitude and 15 ms faster delay. Response in pFC also is similar to the regular protocol (cf. Fig. 3). These results are consistent with AuCPB not being critical to the deviant response in the model, unlike AuRPB.

### 3.5 Feedback error signals

The experiments (Obara et. al. 2023) examined feedback error signals back to the auditory core, finding a class of AuRPB neurons making a strong contribution to the deviant response. Analysis of the marmoset connectivity data shows (Pailthorpe 2023a) that the strong AuRPB (NM #9) link back to the auditory core is to AuRT (NM #3), with weight 2084, while the feedback weight to AuA1 is weak (weight 5). Within the core, AuA1 drives AuRT with a strong link, of weight 2774, while AuRT feeds back to AuA1 with a medium link weight, 430. AuRPB was not an injection site in tracer experiments (Majka et. al. 2016), thus it’s in-link strengths are not known from the experiments; they were estimated here based on statistical trends of out links from the other areas. That suggests it receives strong inputs from the core (AuA1), belt (AL, ML, CM) and from parabelt (CPB), consistent with its key role in auditory response. The strongest outputs from AuRPB, based on the connectivity data, are links back to the core (to AuRT, weight 2.1k) and belt (AuCL 1.0k). Overall, this suggests network effects mediate feedback to the core via multiple links.

Simulations of the regular deviant responses in core, belt and parabelt areas, shown in Figs. 13-15, make explicit the multiple contributions to LFP-Aud. Similarly simulation output allows calculation of individual inputs into AuA1 from linked nodes: these are the driving forces in the DEs describing the NMs (cf. eq. 1-4 of Pailthorpe 2023b). The overall linked driving force into AuA1 is plotted in Supplm. Fig. 8 and is dominated (60 %) by output from AuML, which has the strongest links into AuA1 within the auditory cluster. By comparison AuRPB contributes 0.5% of the driving force into AuA1, suggesting a relatively low direct contribution. AuCPB contributes 16 % of the drive. Feedback from the key output node, TPO contributes 2.5 %. Additional details are discussed in the Suppl. Information at 5. Feedback to AuA1. Overall, these results suggest that feedback from AuRPB to AuA1 is carried over multiple local network paths, rather than only via a direct linkage. The key role of AuRPB identified by experiments likely also reflects specific local circuits within the neural masses not captured by NM models.

### 3.6 Sensitivity tests

A number of implicit assumptions in the model were tested, as discussed in: Supplm.Info. 6. Effects of signal velocity; S.I 7. effect of 2-step links; S. I 8 effects of random number generator; and S.I 9 oddball paradigm stimulus protocol. These extra results confirm the basic relevance of the simulations to the experimental protocols.

## 4. Discussion

Simulations of a neural mass model of the auditory cortex produce NMM deviant response delays comparable to those observed experimentally. Downstream response in pFC were also simulated given that it is a likely target destination of the response signals. A combined model, including pFC areas shows longer response delays due to the signal transmission times. TPO is a key output node of the auditory cluster, with strong in-links from most of the auditory areas. Its strongest out link to A32V, a key in-hub in the marmoset cortex network. This suggests that TPO warrants further investigation in the NMM response. A slower, weaker NMM response is observed in the pFC sub cluster, consistent with such observations in macaque (Camalier et. al. 2019).

The simulations provide detailed information on the responses in individual areas. For instance, sustained beta/gamma bursting, in response to the deviant pulse stimulus to high f areas, is evident in key auditory and hub nodes.

Inactivation of AuRPB induces reduced responses, consistent with observations of its key role in mediating the NMM response; and feedback to the auditory core is shown to be distributed via parallel pathways due to the local network connectivity.

Phase alignment is important when stimulating an oscillator: in-phase stimuli align with and reinforce the oscillations whereas out of phase stimuli work against the oscillator excursions. pFC area A11 has a similar natural frequency to TPO, so more naturally maintains its phase alignment to the forcing signals, while A32V naturally oscillates in the theta band, so drifts in and out of phase with TPO outputs as successive stimuli arrive.

Overall simulation of heterogeneous neural masses, interlinked by the experimentally available link weights, yields negative mismatch deviance delays comparable to those observed experimentally.

### 4.1 Conclusion

A simple, heterogeneous neural mass model, comprising linear oscillators with experimental link weights, is used to describe local clusters in auditory areas and pFC. Nonlinearities enter via the interactions of links neural masses. The model reproduces the observed delay in deviant response to stimuli applied to auditory areas. This suggests that the delay is related to superposition of multiple oscillations and signalling delays. The simulated amplitudes are larger and more persistent than observations, exposing limitations of the model. Likely each area contains multiple cortical columns and sub-circuits with a distribution of characteristics and oscillatory frequencies that superimpose to smooth out the responses. That would require a more sophisticated model than considered here.

## Supplementary Information

### 1. Stimulus Function

The shape of the stimulus function had an effect on high frequency responses, so needed to be controlled for since beta and gamma bursts have been associated with relevant activity (Lundqvist et. al. 2018). A simple square or ramped pulse function (as used in auditory stimulus experiments) has intrinsic high frequency components (evident in fft) due to discontinuities in the function itself and/or its slope. The experimental ramped auditory stimulus (Komatsu et. al. 2015) had 7 ms rise and fall times with a 50ms width of the constant section, giving a total width of 74 ms – that was explored herein. Three pulse shapes were studied, each with that width and a 0.55 s repletion time (also consistent with experiments). Fast Fourier Transform (Matlab fft) results for each yielded fraction of power spectral density in the standard frequency bands: approx. 50% in the delta, 30% theta and ∼5- 10% alpha and beta bands. However they differed in gamma power: 4% for a square pulse stimulus function, 2% for a ramped pulse, and 0.2% for a smooth (sinusoidal) function. To avoid high frequency artefacts induced by the shape of the stimulus, only a smooth pulse function was used hereafter. Care was taken to keep a constant energy input to the NM oscillators so that results are comparable – this was achieved by keeping the integral under the stimulus function constant. A balance of 187 Hz maximum amplitude x (70ms +14) ms width produced stable results, and keeps the dynamics near known bifurcations, and critical transitions (ref), for the models. Use of 50 ms pulse width, as in the experiments, required a stronger peak stimulus, possibly leading to unstable results – yet to be explored. The construction of a single stimulus, comprising a pulse function as typically used in WC/JR simulations, is shown in Supplementary Figure 1. Here the deviant and standard pulses deliver the same energy input to the oscillators; 10 such pulses were applied. Each pulse function delivers 10 “unit pulses” of energy to the oscillator system (being the integral under the pulse curve).

**Supplementary Figure 1.**
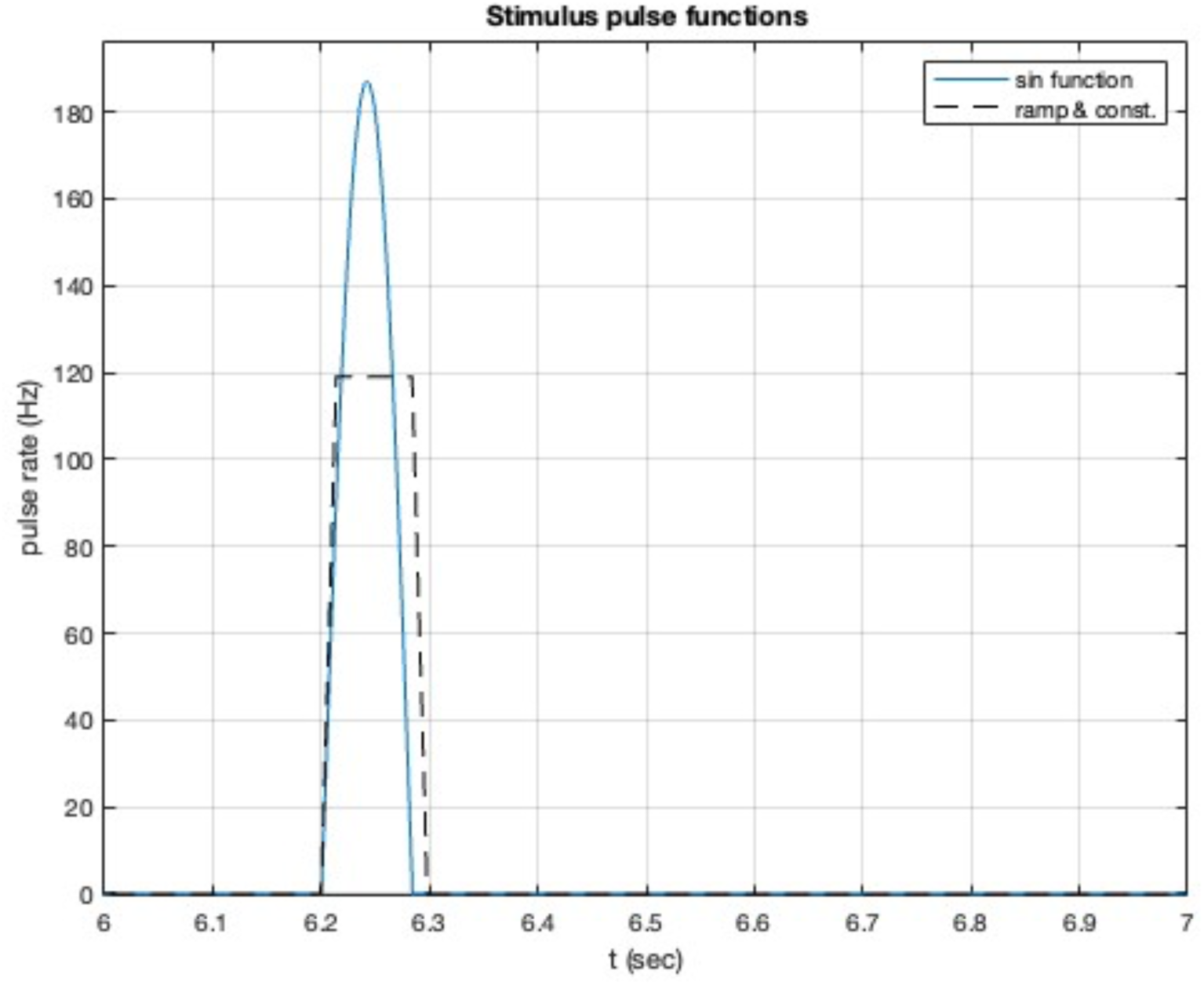
Components of the stimulus pulse function used in the simulations. The ramped form used in the experiments [Komatsu et al 2015] shown as broken black line, while the smooth pulse function used herein is shown in blue. Both functions pulse delivers the same energy input to the NM oscillators.

The experimental protocol (Komatsu et. al. 2015) applied a sound tone that ramped up in intensity over 7 ms, then was constant for a duration of 64 ms, then ramped down. The sharp edges (discontinuous slope) of this functional form can induce spurious high frequency components into the simulation, confounding possible gamma responses, thus was not suitable for a simulation study. Here an equivalent smooth sinusoidal function was constructed to have a similar duration (width) and equal energy input. That was required so that the NM oscillators had a comparable response. The latter was calculated as the integral under the curve, and was equal to 10 unit pulses (units Hz x sec) for both curves shown in Supplm. Fig. 1. Thus 10 pulses contribute 100 unit pulses to the NM oscillators. The scale was chosen to be around 100 Hz, being near bifurcation boundaries known from NM studies. Multiple pulses are repeated at 0.55 s, as in the experiments. The so-called oddball paradigm (Obara et. al. 2023), in which the deviant pulse is 100ms wide, is discussed below.

### 2. Individual deviant responses in Auditory areas

Short time deviant responses are presented in the main text, for comparison with available experimental data; here additional details available from the simulations are presented. Following the initial delay discussed in the main text individual NMs exhibit distinctive structured long time behaviour, as displayed below. Network analysis indicates that TPO is a key output area from the auditory cluster. Its responses were presented in Fig. 4. Further details are shown below. Following the deviant stimulus (5^th^ pulse at 6.2 s) the 6^th^ standard pulse (at 6.75 s) takes effect and induces a new oscillatory pattern with a deviant difference delay (here 125 ms, vs. 129 ms for the deviant stimulus). After the 10 stimulus pulses the deviant difference relaxes and then adopts a different pattern as shown in Supplementary Figures 2 and 3.

**Supplementary Figure 2.**
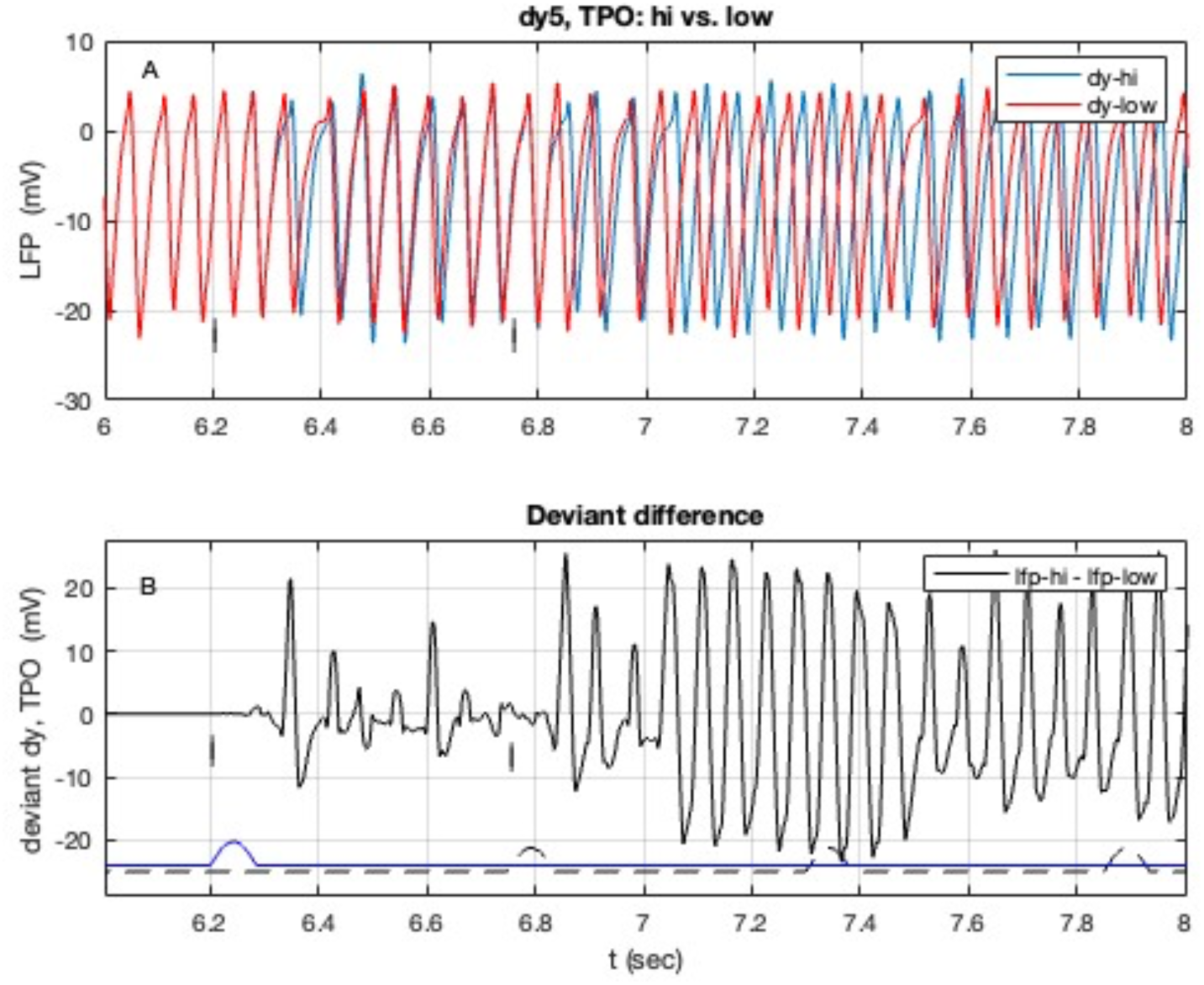
Longer time responses in Auditory area TPO (cf. Fig. 4). **A**) individual responses to stimuli to standard (“low”) and deviant (“hi”) stimuli; **B**) the deviant difference in responses shown in A. Details as in Fig. 2.

**Supplementary Figure 3.**
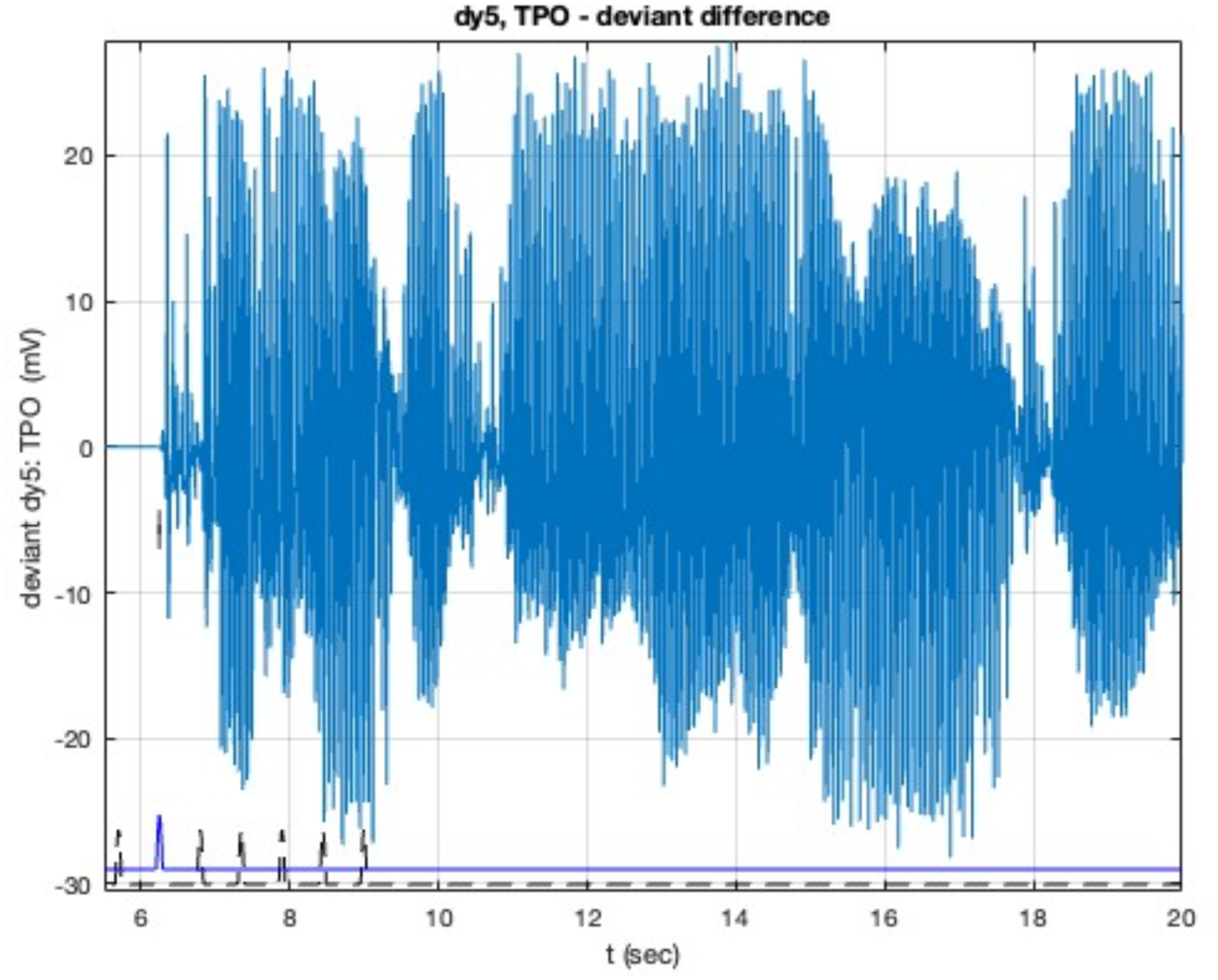
Deviant Δdy response in Auditory area TPO, calculated as high - low responses to stimuli to standard and deviant stimuli. Stimulus pulses shown at bottom. The 5^th^ deviant pulse (to high f areas) is shown in blue, starting at 6.2s (vertical mark).

Given the experimental investigation of area AuRPB, its response is plotted in Supplementary Figure 4, and discussed later.

**Supplementary Figure 4.**
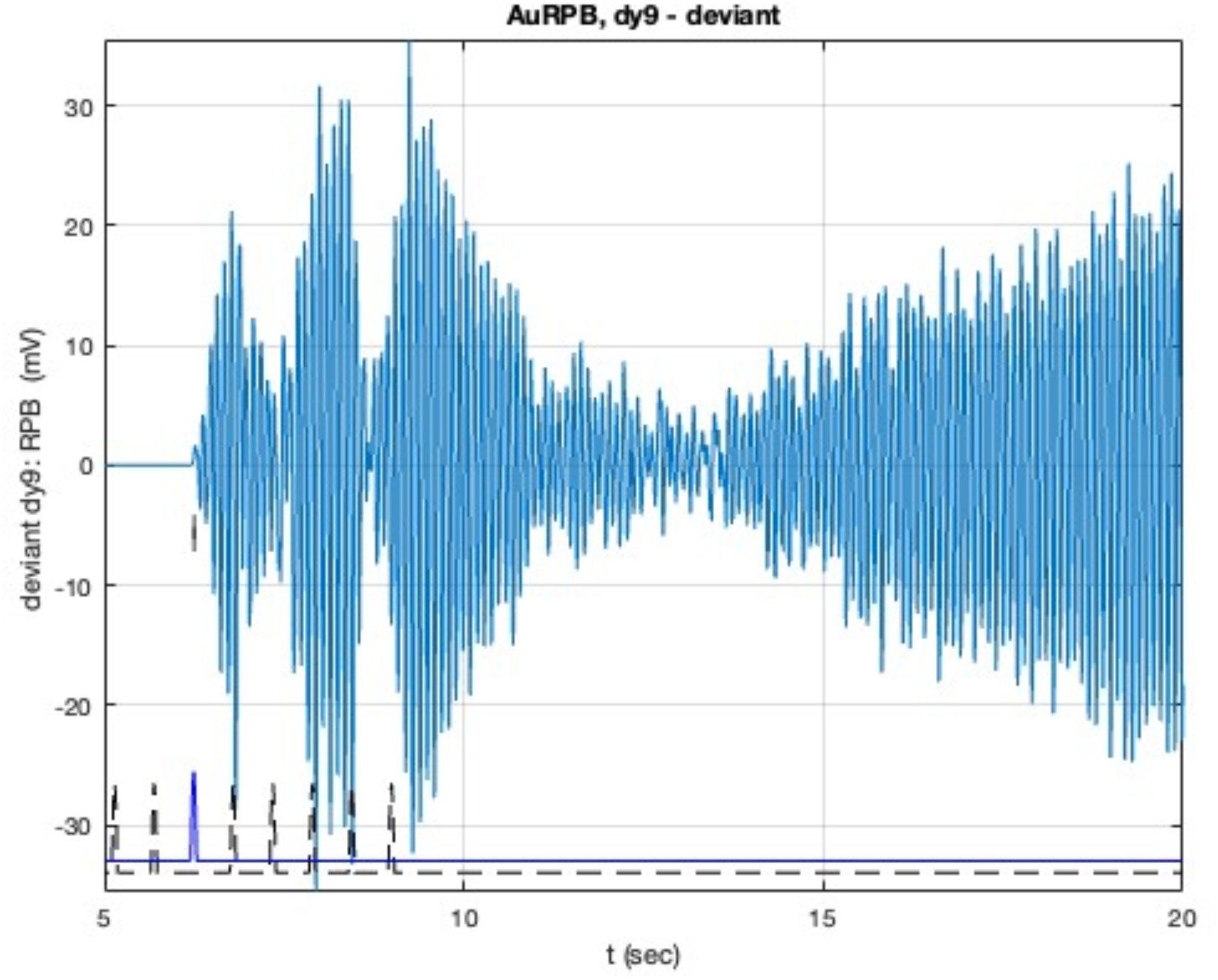
Long time behaviour of deviant response, Δdy in auditory area AuRPB, calculated as high -low responses to stimuli to high and low frequency responsive areas. The 5^th^ deviant pulse (to high f areas) is shown in blue, starting at 6.2s (marked). Stimulus pulses shown at bottom, as in Fig. 2.

### 3. Origin of fast bursts in Auditory areas

Three auditory areas were modelled by 5 sub-population NMs and so can produce gamma bursts in response to a constant stimulus. These areas are explored below in view of the fast bursting induced in pFC (cf. Figs 9 - 11). The response of area AuA1 is shown in Supplm. Fig. 5, and AuCPB in Supplm. Fig. 6. It is evident that AuA1 only oscillates in response to stimuli, showing essentially noise driven fluctuations otherwise. It is evident that the first 4 standard pulses to AuA1 induce only small potential spikes that die out, while the 5^th^ deviant pulse induces sustained fast oscillations – these are in groups of 3 with a period of 50 ms, corresponding to a 20 Hz oscillations; the groups repeat at 175 ms or 5.7 Hz. For CPB the standard pulses each induce 2 oscillations that rapidly decay, while the deviant pulse induces 4 double bursts in with a period of 67 ms, corresponding to a 14.9 Hz oscillation. Area AuRT is also in this group: it is tightly coupled into the auditory cluster so was included in the simulations. Its response is in Supplm. Fig. 7. Standard pulses induce sustained bursts with a period of 50 ms, corresponding to 20 Hz oscillations, along with occasional faster large spikes at 42 ms, or 23.8 Hz. The 5^th^ deviant pulse does not induce such a spike. For this NM the oscillations continue after the stimuli cease.

**Supplementary Figure 5.**
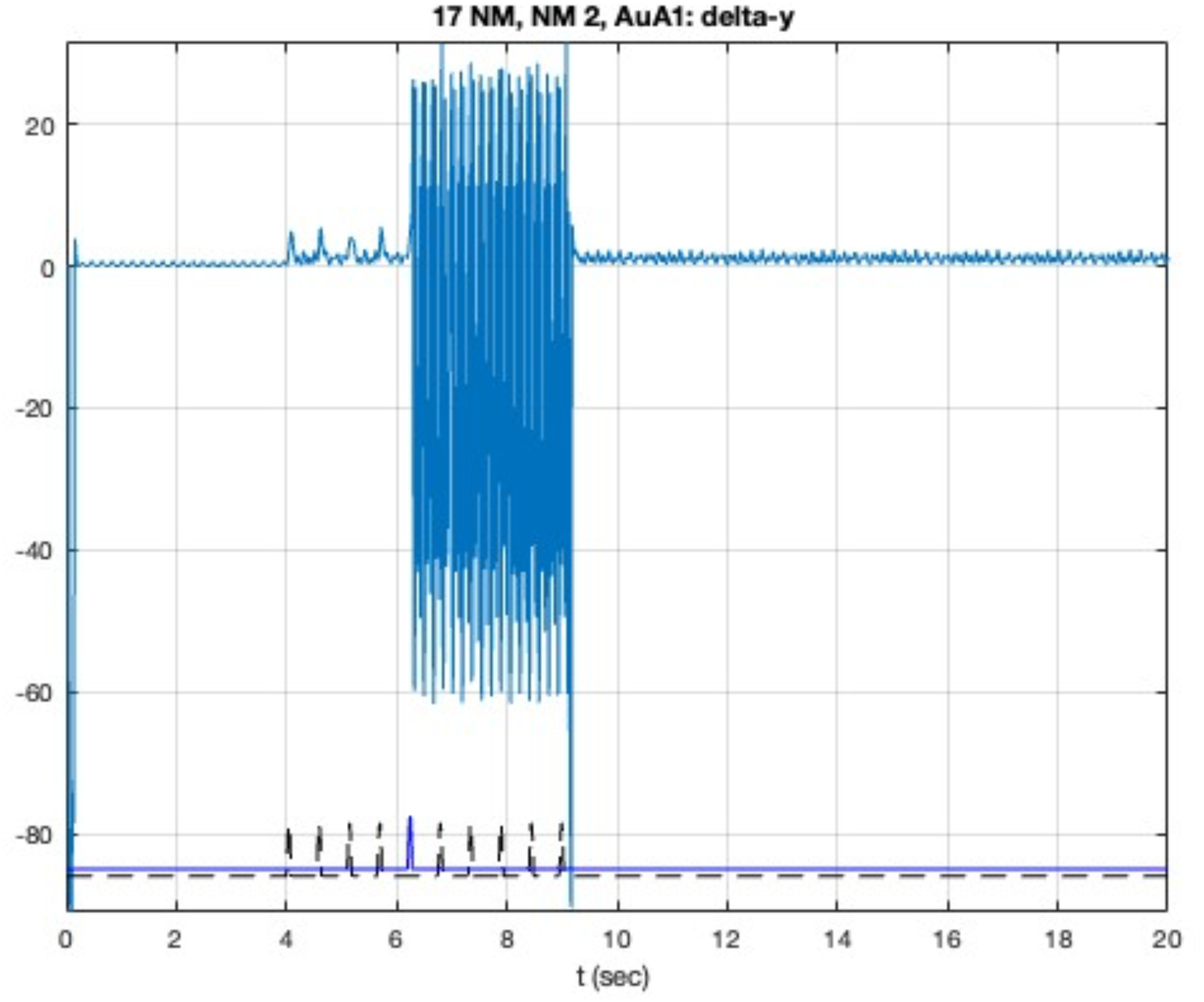
The response dy in auditory area AuA1 to the deviant stimulus (9 pulse to low and one pulse to hi f areas). Stimulus pulses shown at bottom, as in Fig. 2. The 5^th^ deviant pulse (to high f areas) is shown offset in blue, starting at 6.2s (vertical mark).

**Supplementary Figure 6.**
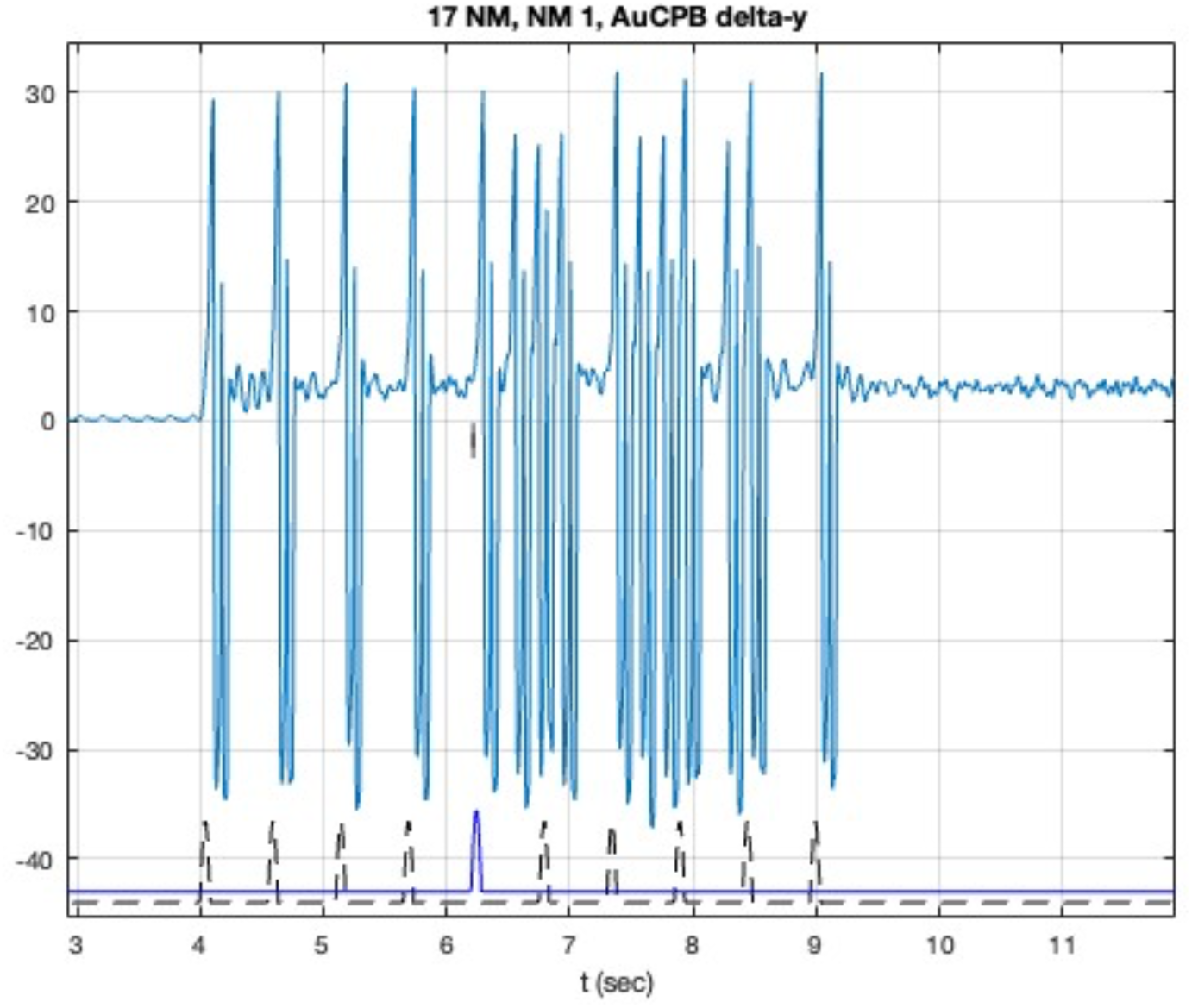
Bursting voltage response dy, in auditory area AuCPB to the deviant stimulus (9 pulse to low and one pulse to hi f areas). Stimulus pulses shown at bottom, as in Fig. 2. The 5^th^ deviant pulse (to high f areas) is shown offset in blue, starting at 6.2s (vertical mark).

**Supplementary Figure 7.**
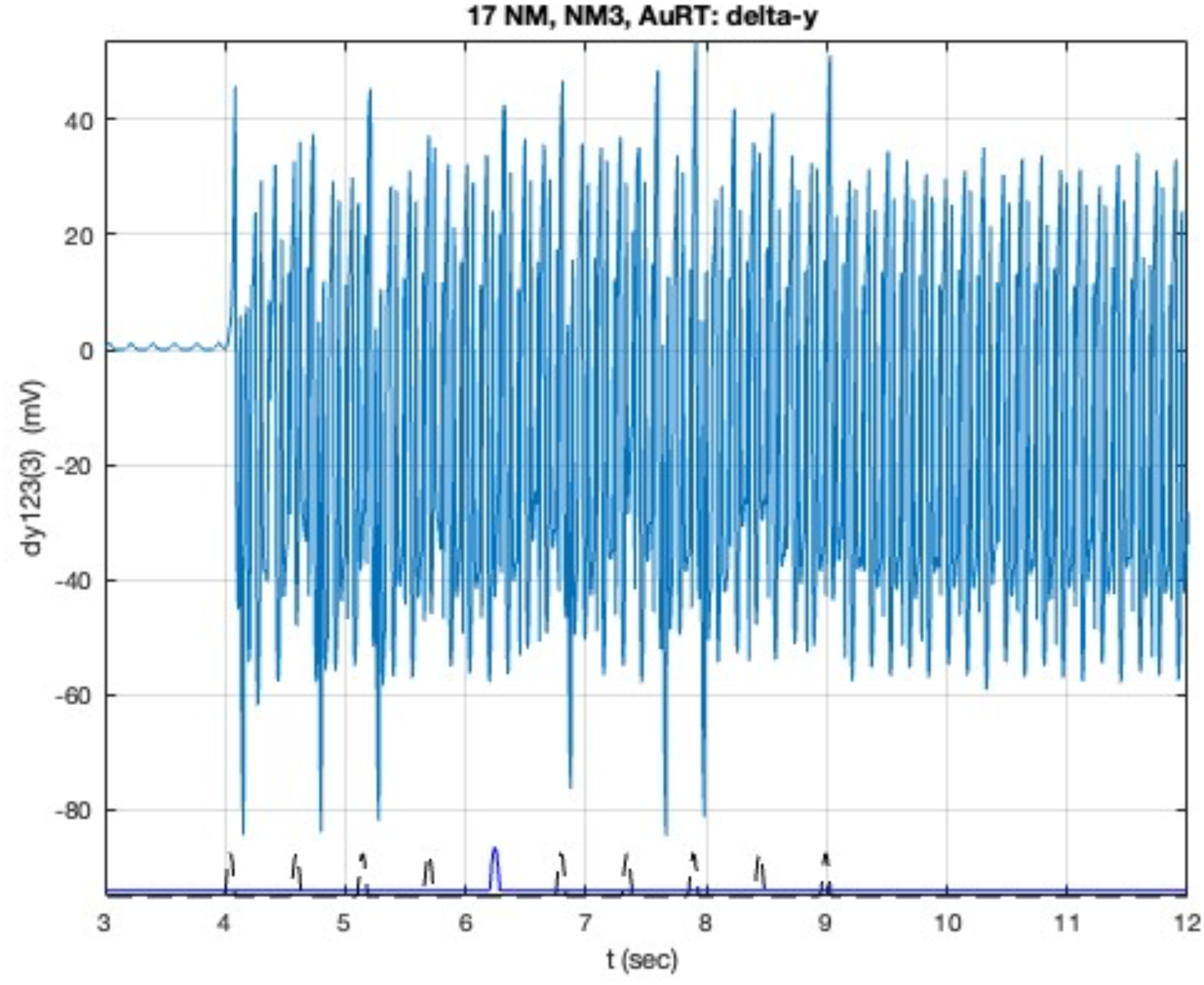
The deviant response, dy in auditory area AuRT to the deviant stimulus (9 pulses to low and one pulse to hi f areas). Stimulus pulses shown at bottom, as in Fig. 2. The 5^th^ deviant pulse (to high f areas) is shown offset in blue, starting at 6.2s (vertical mark).

### 4. Role of RPB

This was discussed in 3.4 Methods and 3.5 Results; further details of deviant responses in specific auditory areas are presented here. The waveform for AuA1 is relatively unchanged so is not plotted here. TPO is the key output node from the auditory cluster so its waveform is plotted in Supplm. Figure 8.

**Supplementary Figure 8.**
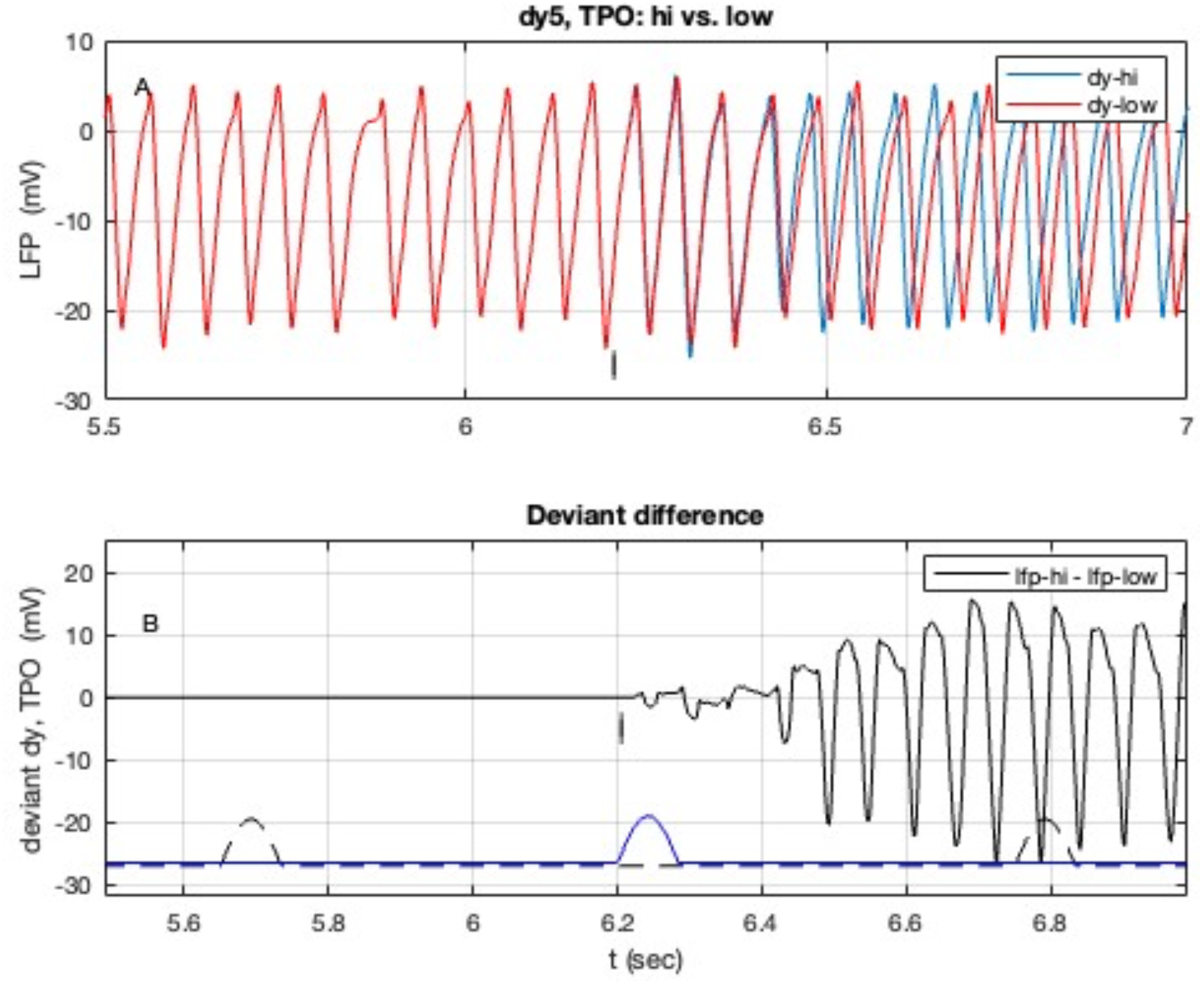
Components of the deviant response in auditory area TPO with AuRPB inactivated. **A**) individual responses to stimuli to high and low frequency responsive areas; **B**) the difference in responses. Details as in Fig. 2.

### 5. Feedback to AuA1

(cf. related discussion in main text at 3.6 Feedback error signal)

The DEs describing the NMs (cf. eq. 1-4 of P 2023b) allow identification of the external driving forces causing the voltage oscillations. The right hand side of those equations shows that this is the sum of (A/τ_e_) S[dyj(t)] (units mV/s^2^) for each linked node j. That sum of inputs to AuA1 is plotted in Supplm. Figure 9.

**Supplementary Figure 9.**
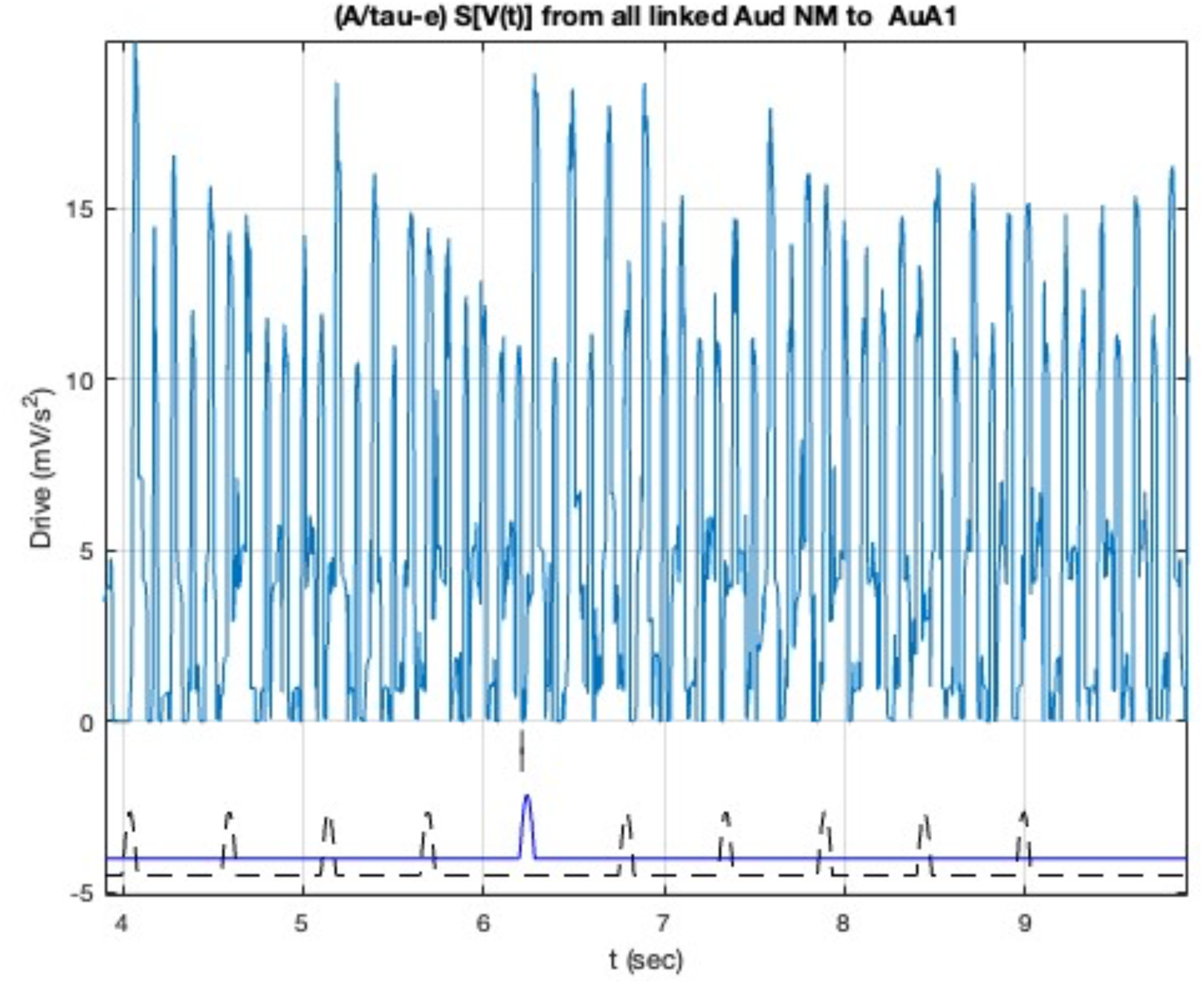
Total driving force at AuA1 from linked auditory NMs, in the deviant stimulus protocol. Stimulus pulses plotted at the bottom; details as in Fig. 2.

The 5^th^, deviant, stimulus pulse causes a prompt (delay 85 ms) increase in drive, resulting in the oscillatory response shown in Fig 6A. Note that other local inputs from slow and fast inhibitory sub-populations also contribute, possibly suppressing oscillations. This total linked driving force into AuA1 is dominated (60 %) by output from AuML, which has the strongest links into AuA1 within the auditory cluster. By comparison AuRPB’s contribution peaks at 0.09 mV/s^2^, or 0.5% suggesting a relatively low direct contribution. From the belt, AuML contributes 60 %, AuCL 22 %, AuAL 6 % and AuCM 6 %. Locally, in the core, AuRT contributes 0.5 % and AuCPB contributes 16 % of the drive. Feedback from the key output node, TPO contributes 2.5 %. These results suggesting that feedback from AuRPB to AuA1 is carried over multiple local network pathways.

### 6. Sensitivity test: phase alignment and axonal signal velocity

A simple test of the role of phase alignment is: setting the long range (axonal) signal velocity to 1 m/s lengthens the delay in the deviant stimulus reaching pFC: eg. for the dominant, direct TPO – A32V path the delay increase from 1.3 ms to 13.4 ms. While for the 2-step paths via multiple waypoints, the average (across relevant waypoints) delay increases from 2 ms to 19.9 ms. For A32V the dominant frequency is 5.06 Hz, or period 198 ms; thus the change in delay is 18% of the positive (half) cycle of dy at A32V – a significant phase shift. The two simulations produce identical wave forms at Auditory areas, and similar ones in pFC. In response to those changes the deviant time (to the first negative minimum voltage difference (cf. Fig. 2) of ΔLFP-pFC increases from 148 to 186 ms. That 38 ms shift in delay can be compared with the 17.8 ms increase in signal transmission delay. The pulsing gamma bursts (at 32.3 Hz) from A10 now have a new form, shown in Supplm. Figure 10. Compared to the regular result in Fig. 12 the ringing gamma bursts have the same general form but here have slightly larger amplitude and a faster beating pattern. This illustrates one effect that changing signal phase has on signal processing within pFC.

**Supplementary Figure 10.**
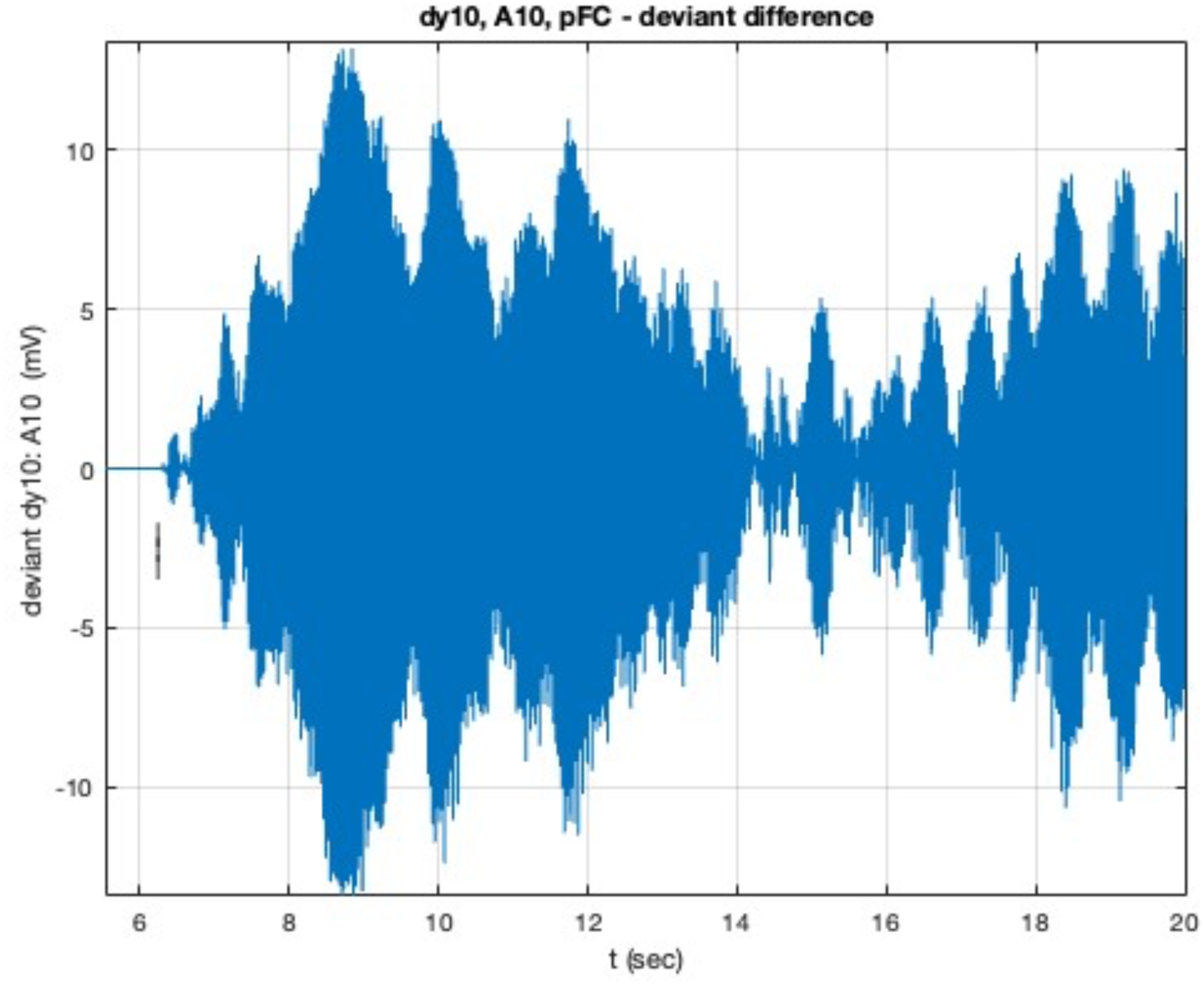
Deviant difference Δdy response in pFC area A10 with long range signal velocity set to 1 m/s, calculated as deviant - standard responses to stimuli. The 5^th^ deviant pulse (to high f sensitive areas) starts at 6.2s (vertical mark). Stimulus pulses not shown.

This test also sheds light on an implicit assumption in the present model. The local links (≌ 1.5 - 6 mm length) in each cluster have signal delays due to an assumed 1 m/s local signal velocity. That also applied to the direct feedforward and feedback links between the two clusters, despite those links being about twice as long (≌ 12-16 mm). That assumption was implicit in the algorithm to solve the JR equations. The general form of responses is preserved, with details slightly changed as illustrated by comparing Fig. 12 and Supplementary Figure 10.

### 7. Role of direct and indirect auditory - pFC links

The simulation results in the main text included the effect of 2-step links via multiple waypoints. Here that assumption is probed by incorporating only the direct links between the auditory and pFC clusters. The long range signal velocity was reset to 10 m/s, as in the main text. The first delay in ΔLFP-Aud is 117 ms compared to 132 ms (cf. Fig. 2B); and for ΔLFP-pFC it is 137 ms vs. 145 ms. The faster responses are consistent with the shorter signal travel times associated with the direct links, compared to the 2-steps links used in the main text. In each case there are two ΔLFP dips in rapid succession (beta band oscillation), in contrast to the main results with only one dip. Another illustrative result is the deviant difference response Δdy-A10 in pFC using only direct links between Aud – pFC, as shown in Supplm. Figure 11. The structured rapid bursting after the deviant pulse evident in Fig. 12 is absent here, replaced by a sustained oscillation, that grows in amplitude.

**Supplementary Figure 11.**
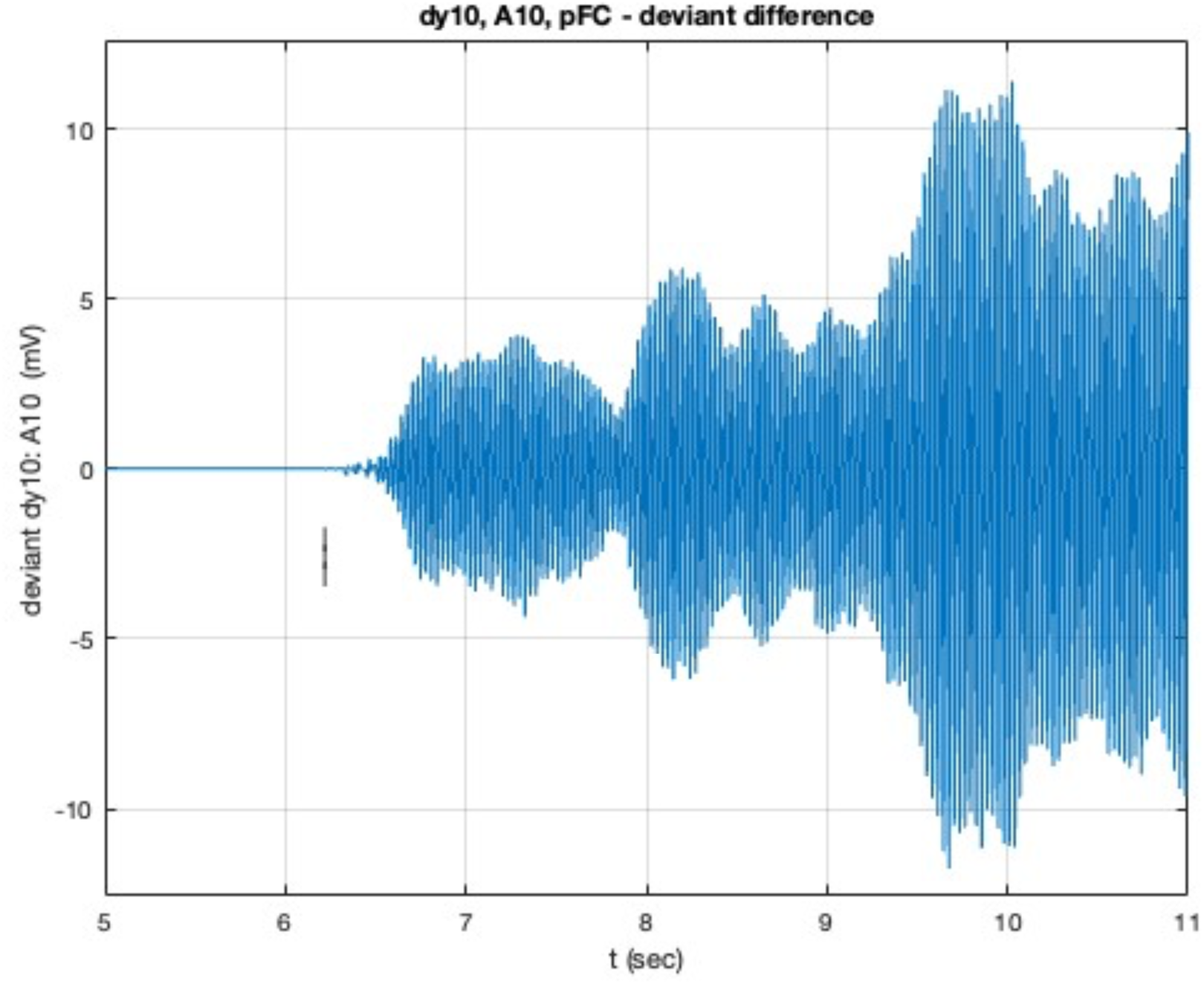
Deviant difference, Δdy, response in pFC area A10 with direct Aud – pFC links only, calculated as high -low responses to stimuli to high and low frequency responsive areas. The 5^th^ deviant pulse starts at 6.2s (vertical mark). Stimulus pulses not shown.

### 8. Effects of random number generator

The deviant response is a small difference between two larger quantities, so is sensitive to numerical methods. As is usual in NM simulations a random pulse input (here 1 sd gaussian random noise) is used to simulate general inputs from surrounding nervous tissues. In the standard NMM experiment (Kamatsu et. al. 2015; Obara et. al. 2023) the deviant stimulus is contemporaneous with the standard stimulus, so the bath of surrounding inputs should be similar, if not identical. To simulate that the simulation with the deviant pulse train (9 low + 1 hi f pulse[s]) should use the same random number stream as the standard stimulus (10 pulses to low f areas). An illustration of the effect is the response of LFP for auditory areas, as shown in Supplm. Figures 12A (a repeat of Fig. 2), with the same random number stream, and 12B where the random number generator is reset (via the system clock) prior to the deviant stimulus protocol. In the latter case the different noise inputs cause a difference in LFP responses due to the same standard pulses, even prior to the 5^th^ deviant pulse. The NMM delay is still evident, at 129 ms (Supplm. Fig. 12B) compared to 132 ms (Supplm. Fig 12A) with randon number generator reset. The gamma bursting patterns are similar overall but differ in details. The NMM dip and delay are comparable for auditory areas, while for pFC such a dip is harder to discern, being of a similar magnitude to noise induced fluctuations prior to the onset of the 5^th^ deviant pulse.

**Supplementary Figure 12.**
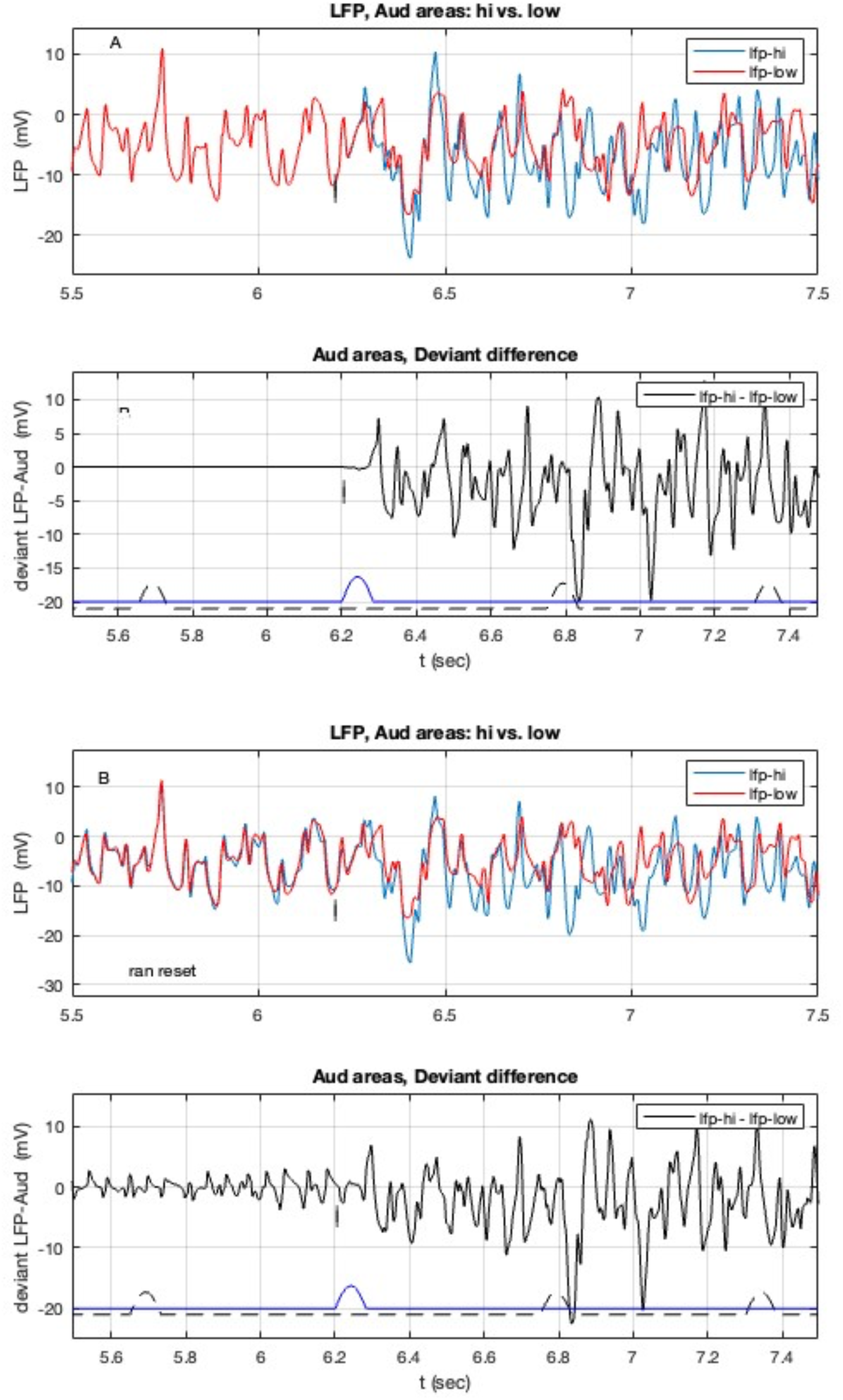
Deviant difference ΔLFP-Aud response for **A**) The same random number stream was used for both standard and deviant simulations; top: individual waveforms and bottom their deviant difference **B**) the random number generator was reset prior to the deviant simulation. Standard stimulus pulses shown as black broken lines. The replacement 5^th^ deviant pulse (to high f areas) is shown offset in blue, starting at 6.2s (vertical marks).

### 9. Oddball paradigm

A related experimental protocol was the so-called oddball paradigm (Tani et. al 2018; Obara et. al. 2023) in which 50 ms duration tones, with 7ms rise and fall times, were applied as the standard stimulus. The deviant replaced one tone pulse with a 100 ms duration tone of the same frequency. The protocol adopted herein, illustrated in Fig. 1, is slightly different and simpler – in particular all pulses deliver the same energy input to the NMs. Now the standard 70+14 ms wide smooth pulses (cf. Supplm. Fig. 1) were replaced by 50+14 and 100+14 ms pulses, respectively. Now a 100 ms long 5^th^ deviant stimulus is applied to high f areas. Thus the standard pulse delivers 7.6 unit pulses of energy (vs. 10 in the main text), while the deviant delivers 13.6 unit pulses. It is clear that the deviant stimulus here delivers more energy compared to the results presented in the main text. The LPP-Aud deviant response is shown in Supplm. Figure 13. Now the delay in deviant response, ΔLFP-Aud, was 131 ms, compared to 132 ms (cf. main text and Fig. 2) for auditory areas. With more energy delivered by the deviant stimulus pulse the amplitude of the initial deviant difference is larger. For pFC the delay is 149 ms vs. 145 ms in Fig. 2

**Supplementary Figure 13.**
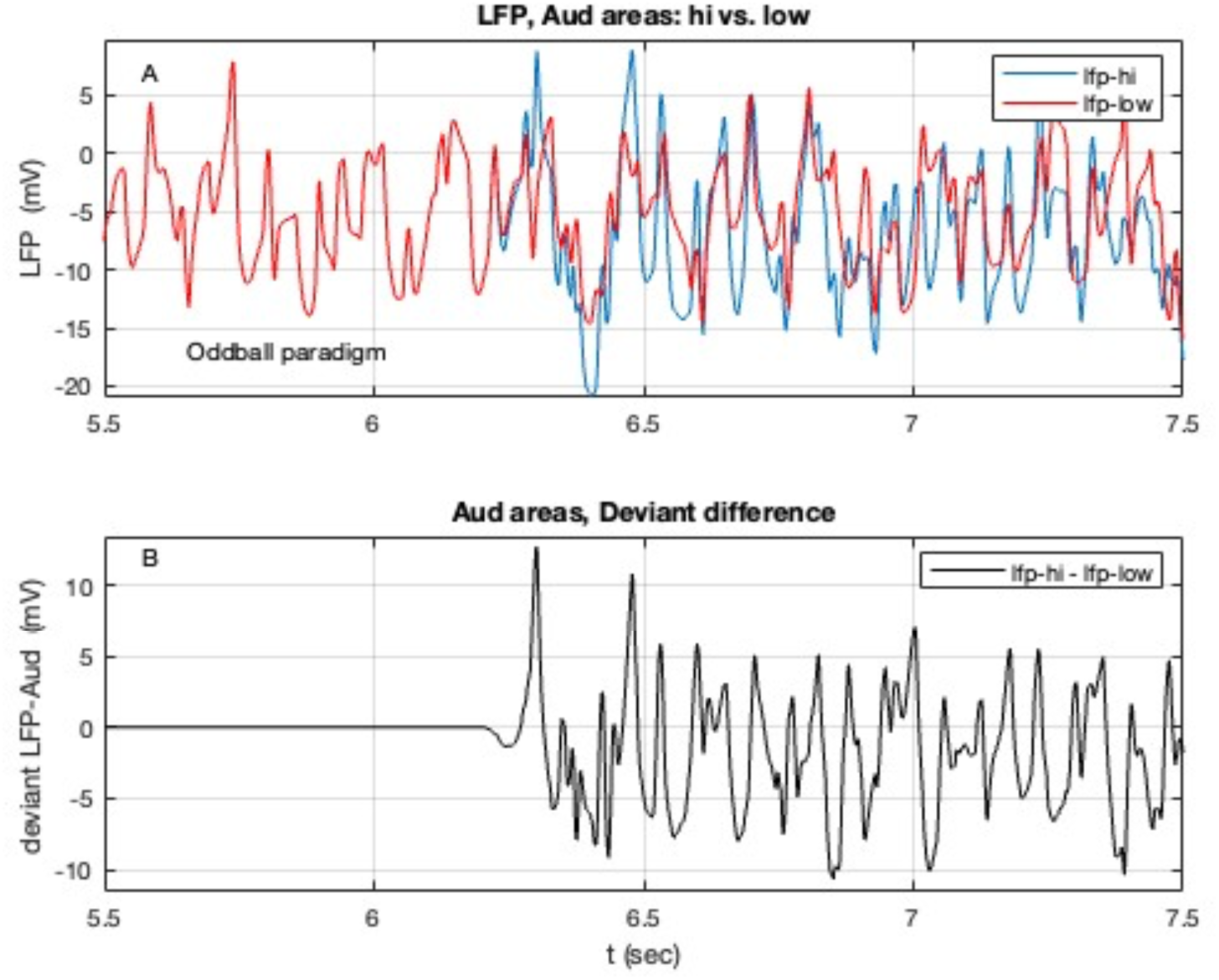
LFP-Aud response in the oddball paradigm for **A**) individual waveforms (standard and deviant stimuli) and **B**) their deviant difference ΔLFP-Aud. Details as in Supplm. Fig. 11.

In each case a structured gamma bursting signal follows the deviant stimulus, particularly in the identified pFC hub for out links, A10 (cf. Fig. 12). With the longer deviant pulse delivering more energy the initial (6.2 – 7.5 s) bursts are larger amplitude and then revert to the regular pattern at longer times (cf. Fig 12).

## Notes

### Competing Interest Statement

The authors have declared no competing interest.

